# yFibronectin-Based Nanomechanical Biosensors to Map 3D Strains in Live Cells and Tissues

**DOI:** 10.1101/2020.02.11.943696

**Authors:** Daniel J. Shiwarski, Joshua W. Tashman, Alkiviadis Tsamis, Jacqueline M. Bliley, Malachi A. Blundon, Edgar Aranda-Michel, Quentin Jallerat, John M. Szymanski, Brooke M. McCartney, Adam. W. Feinberg

## Abstract

Mechanical forces are integral to a wide range of cellular processes including migration, differentiation and tissue morphogenesis; however, it has proved challenging to directly measure strain at high spatial resolution and with minimal tissue perturbation. Here, we fabricated, calibrated, and tested a fibronectin (FN)-based nanomechanical biosensor (NMBS) that can be applied to cells and tissues to measure the magnitude, direction, and dynamics of strain from subcellular to tissue length-scales. The NMBS is a fluorescently-labeled, ultrathin square lattice FN mesh with spatial resolution tailored by adjusting the width and spacing of the lattice fibers from 2-100 µm. Time-lapse 3D confocal imaging of the NMBS demonstrated strain tracking in 2D and 3D following mechanical deformation of known materials and was validated with finite element modeling. Imaging and 3D analysis of the NMBS applied to single cells, cell monolayers, and Drosophila ovarioles demonstrated the ability to dynamically track microscopic tensile and compressive strains in various biological applications with minimal tissue perturbation. This fabrication and analysis platform serves as a novel tool for studying cells, tissues, and more complex systems where forces guide structure and function.

## Introduction

Cell-generated mechanical forces transmitted via cell-cell and cell-extracellular matrix (ECM) interactions are integral to a wide range of processes from cell migration^1–6^ and contractility^7^ to tissue morphogenesis^8–14^ and regeneration^15–17^. Many diseases states are also characterized by impaired force generation such as dilated cardiomyopathy^18^, or changes in ECM mechanical properties that in turn alter mechanosensitive gene expression such as in fibrosis^19^. Yet in most cases the underlying mechanobiology is not well understood, in part because direct measurement of forces in living tissue during dynamic processes has proved difficult. Further, cells and tissues are spatially heterogeneous, with properties that can very over the span of a few micrometers up to centimeters and forces generated on the cellular scale that combine to produce the macroscale forces that shape tissue formation.^20,21^ A major challenge in the field is to directly map these cellular and tissue-scale mechanical forces in 3D in both *in vitro* and *in vivo* systems. To do this, we need to develop new measurement tools that can span multiple length scales to enable tracking while minimally perturbing the biology of interest. However, forces do not need to be measured directly, and instead deformation has been and continues to be used widely to determine strain and serve as a proxy to determine stress and related material and biomechanical properties.

There have been a number of important advances to measure strain in cells and tissues using a combination of computational, optical and combined methods. Researchers have measured strains using finite element modeling and related computational techniques such as digital image correlation and finite element analysis (FEA) in order to determine strain fields and estimate stresses based on known or approximated material properties.^22,23,25,26^ Advanced optical techniques include live cell imaging-based optical sensors (Förster resonance energy transfer, FRET),^24^ fluorescence based oil microdroplets,^21^ mechanical traction sensors in 2D and 3D determine traction forces.),^25–27^ birefringence,^28^ and force inference.^29^ Together, these methods have advanced the field of biomechanics and mechanobiology, and provided new insights into the role of biological forces in cell and tissue development and function. However, experimental limitations such as cell and tissue perturbation, limited field-of-view, sparse measurement density, and significant computational overhead, have spurred the continued development of newer approaches. There remains need for a method that can combine (i) direct measurement of strain, (ii) high spatial and temporal resolution, (iii) tracking and mapping in both 2D and 3D, and (iv) minimal perturbation of the cell or tissue system.

Here we have developed a fibronectin (FN) based nanomechanical biosensor (NMBS) that provides the capability to quantify the location, direction, and magnitude of strain on a 3D surface over time, from subcellular to tissue length scales. The NMBS is composed of an ultra-thin mesh (~10 nm thick) of fluorescently-labeled FN that can readily adhere to and integrate onto the surface of cells and tissue. The engineered micropattern of the fluorescent NMBS mesh acts as fiducial markers to track strain dynamically during developmental and physiological processes via live 3D fluorescence imaging. The NMBS is created utilizing an adaptation of our surface-initiated assembly (SIA) technique, which can be used with a range of ECM proteins (e.g., collagen type IV, laminin)^30^ and mesh micropattern designs depending upon experimental requirements.^31^ Prior knowledge of the tissue mechanical properties is not required, and spatial resolution of the NMBS can be tuned by defining the width and spacing of the fibers that make up the mesh from 2 μm to 100 μm spacing, enabling measurements from <1 μm to >1,000 μm. Confocal and multiphoton imaging of NMBS deformation over time is then analyzed using a computational and visualization software package for 3D image segmentation, tracking, and mapping of strain. Validation is achieved using materials of known material properties combined with FEA of deformation fields. Finally, we demonstrate the ability to directly quantify cell-generated 3D tensile and compressive mechanical strains at both cellular and multicellular resolution by applying the FN-based NMBS to the surface of single cells, cell monolayers, and developing tissue.

## Results

### Fabrication of the Fibronectin-based NMBS

The NMBS is a fluorescently-labeled FN square lattice mesh with tunable dimensions that can be applied to cells and tissues and tracked in 3D using time-lapse confocal and multiphoton imaging. The NMBS is fabricated using an adaptation of surface-initiated assembly (SIA), which is a technique to microcontact print ECM proteins onto a thermo-responsive poly(N-isopropylacrylamide) (PIPAAm) surface and then release the ECM proteins (e.g., FN, laminin, collagen type IV, fibrinogen) as an assembled, insoluble network with defined geometry.^31^ Our previous work has demonstrated that SIA can be used to engineer sheets of ECM proteins to shrink wrap cells in a defined ECM niche,^32^ create a basement membrane to enhance endothelial cell growth,^33^ and to micropattern ECM proteins on to microtopographies.^30^ Here, instead of creating an ECM scaffold we have repurposed the SIA approach to form the NMBS. Computer-aided design (CAD) software is used to define the geometry of the mesh, and in this initial work we selected a square-lattice mesh because we can easily change the width and spacing of the fibers, and spaces between fibers, based on the experimental purpose.

The overview of the NMBS fabrication process and application to cells and tissues is illustrated in Figure 1. Briefly, the design of the NMBS is created in CAD, transferred to a transparency photomask, and used to photolithographically pattern a photoresist coated glass wafer (Figure 1a). A polydimethylsiloxane (PDMS) elastomer stamp is formed by casting on the patterned wafer (Figure 1b) and is then inked with fluorescently-labeled FN (Figure 1c). The PDMS stamp is then used to microcontact print the FN onto a PIPAAm coated glass coverslip (Figure 1D) and then removed to create the NMBS on the PIPAAm surface (Figure 1e). To conform to the topology of cells and tissues, the NMBS must be transferred from the rigid glass coverslip to a flexible carrier. To do this a gelatin carrier film is brought in contact with NMBS-patterned PIPAAm-coated coverslip and then room temperature water is added to dissolve the PIPAAm and transfer the NMBS to the gelatin (Figure 1f). The NMBS is then applied to various sample types including hydrogel test strips (Figure 1g), cell monolayers (Figure 1h), and developing tissues (Figure 1i). In each case the NMBS is transferred by incubation at 37°C for 5 min to melt the gelatin, integrating the NMBS onto the desired surface. The NMBS can be tailored over a wide range, but we found that dimensions with line width and spacing of 2 µm x 2 µm, 20 µm x 20 µm, and 100 µm x 100 µm were ideal for measuring strain from the subcellular to multicellular length scales and could be created on the PIPAAm surface and transferred to the gelatin carrier with high fidelity (Figure 1j). Finally, the thickness of the NMBS was ~4 nm by atomic force microscopy (Figure 1k, l), verifying the nanoscale dimension and the formation of an ultra-thin mesh that should minimally perturb the surface it is attached to.

**Figure 1.**
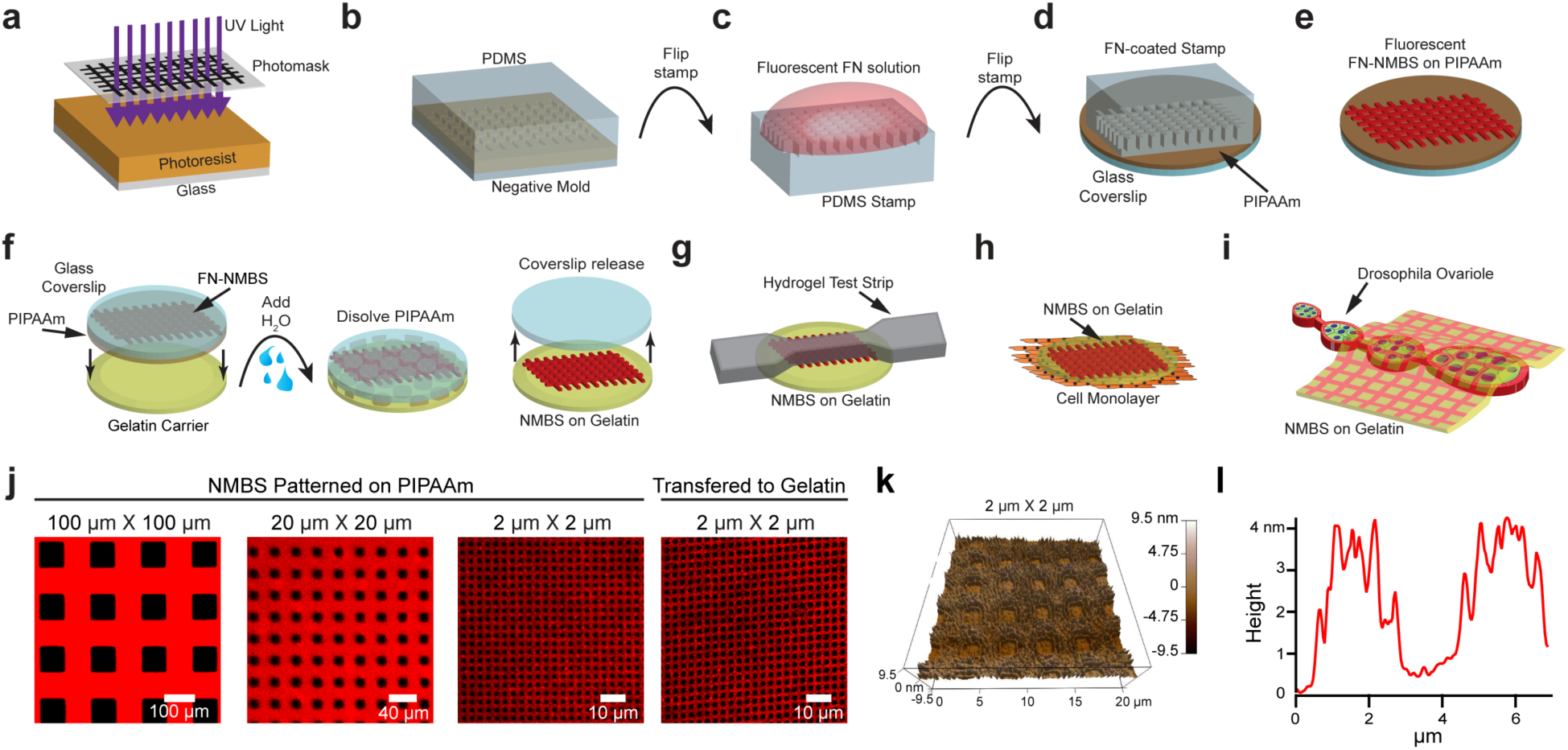
Microfabrication, Experimental Set-up, and Analysis of the NMBS. **a**. Exposure of photoresist-coated glass wafer to ultraviolet (UV) light through a custom mesh photomask. **b**. Casting of polydimethylsiloxane (PDMS) over topographically patterned photoresist-coated glass wafer. **c**. PDMS stamp coated with fluorescently labeled fibronectin (Alexa-555 FN, 50 µg/mL) solution. **d**. FN-coated stamp is microcontact-printed onto a poly(N-isopropylacrylamide) (PIPAAm)-coated glass coverslip. **e**. NMBS is patterned on the sacrificial substrate PIPAAm. **f**. The NMBS-PIPAAm coverslip is placed NMBS-side down onto a gelatin type A carrier. Water is added to dissolve the PIPAAm and transfer the NMBS to the gelatin which releases the glass coverslip. **g, h, i.** The NMBS is applied to dog bone hydrogel test strips (g), the top surface of cell monolayers (h), and the surface of drosophila ovarioles (i) by placing the NMBS-gelatin carrier NMBS-side down on to the sample and raising the temperature to 37°C to melt the gelatin and integrate the NMBS with the sample. **j**. Alexa-647 fibronectin NMBS of varying resolution from large 100 µm X 100 µm to small 2 µm X 2 µm mesh sizes patterned onto PIPPAm and transferred onto gelatin. **k**. AFM height retrace showing the topology of the patterned NMBS. **l.** Line plot from the height retrace reveals an average thickness for the fibronectin NMBS of ~ 4 nm.

### Validation of the NMBS to Track Strain

To validate and benchmark the NMBS as a microscopic strain sensor we performed controlled mechanical testing while visualizing microscopic deformations of the NMBS through fluorescence microscopy. First, a PDMS “dog bone” shaped tensile test strip was designed and molded (Figure S1) and NMBS with 10 µm wide lines and 100 µm was transferred across the gauge length (Figure 1g). This test strip was then mounted in a custom-designed uniaxial testing system that enables simultaneous microscopic imaging of the NMBS and macroscopic imaging of the PDMS (Figure 2a). During tensile testing elongation of the PDMS test strip is measured by an increased distance between the fiducial marks (small black dots on the PDMS in the gauge length, Figure 2b). Fluorescence images of the NMBS pre-strain (Figure 2c) and following 45% uniaxial strain (Figure 2d) show the microscopic deformations tracked by the NMBS mesh. The initial observation state, *l*_0_, is considered as the zero-strain reference configuration. Subsequent deformation of the NMBS, *l*, relative to the initial reference state, *l*_0_, enables calculation of engineering strain as 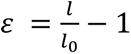 within each NMBS segment. Time-lapse fluorescence imaging of the NMBS provides tracking of both tensile and compressive strain at the microscopic scale based on mesh deformation. Quantification of strain is performed using custom MATLAB software by converting the NMBS fluorescence image into a binary image, identifying intersection nodes, connecting the nodes to form a rectangular mesh, and overlaying it onto the original image (Figure 2e). Node-node distances are measured as segment lengths for calculation of engineering strain to demonstrate tensile (positive) and compressive (negative) strains with respect to their undeformed reference length (Figure 2f). The segments are further color coded according to their strain value to obtain a global map of the tensile and compressive strains (Figure 2f).

**Figure 2.**
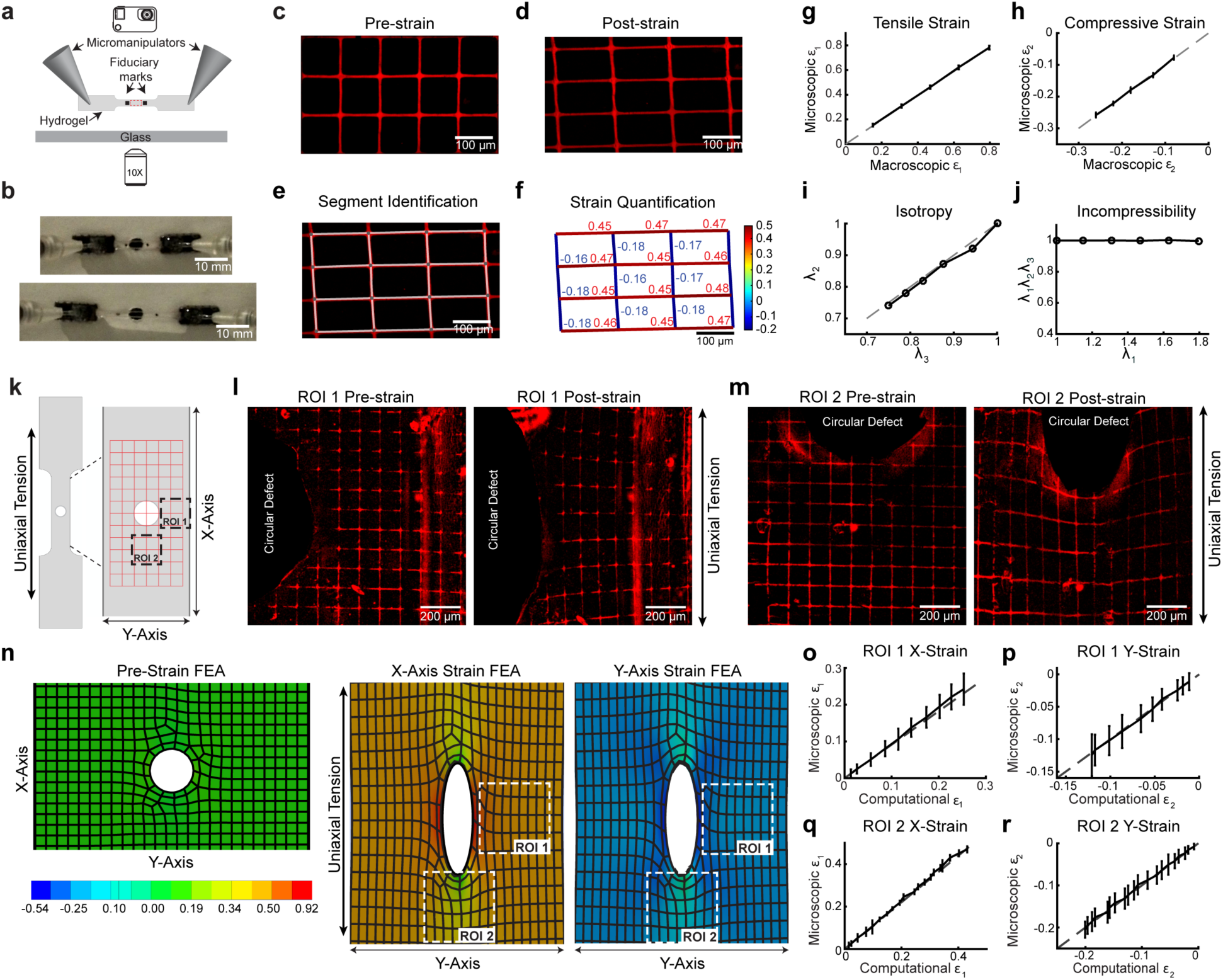
Uniaxial tensile testing and Finite Element Analysis of the NMBS on a PDMS test strip. **a**. A PDMS dog bone hydrogel test strip with applied NMBS is mounted on micromanipulators attached to a wide-field fluorescence microscope to view the NMBS (100 µm X 100 µm X 10 µm) from underneath while tracking fiduciary marks on the top surface during uniaxial tensile testing. **b.** Example images of the PDMS test strip in the pre-strain and post-strain states. **c**. Representative fluorescence image of pre-strained NMBS prior to tensile testing. **d**. Representative fluorescence image of deformed NMBS following tensile testing. **e**. Image segmentation and analysis is performed to identify rectangular mesh NMBS nodes and segments are then overlaid onto the deformed image. **f**. Segment lengths are converted into tensile (positive) and compressive (negative) mechanical strains with respect to their pre-strain reference length. A color map is applied to the mesh segments to reveal locations of tensile and compressive mechanical strains. **g.** The microscopic tensile strain ε_1_ of the NMBS strongly correlates with the macroscopic tensile strain ε_1_ of PDMS (mean ± S.D.). **h**. The microscopic compressive strain ε_2_ of the NMBS strongly correlates with the macroscopic compressive strain ε_2_ of PDMS (mean ± S.D.). **i.** There is a strong correlation between the stretch ratios λ_2_ and λ_3_ confirming that PDMS is an isotropic material. **j.** The product of the stretch ratios λ_1_λ_2_λ_3_ is equal to 1 confirming that PDMS is incompressible. **k.** A circular defect in the PDMS test strip was introduced to track heterogeneous strain fields with the NMBS during uniaxial tensile testing. **l.** Representative fluorescence images from ROI 1 (region adjacent to the circular defect and antiparallel to the uniaxial tension) of the NMBS pre and post tensioning. **m.** Representative fluorescence images from ROI 2 (region adjacent to the circular defect and parallel to the uniaxial tension) of the NMBS pre and post tensioning. **n.** Finite element analysis of a PDMS test strip with a circular defect in the pre-strain, X-strain, and Y-strain states following 40% uniaxial strain. **o.** Quantitative analysis of ROI 1 (region adjacent to the circular defect and antiparallel to the uniaxial tension) reveals a strong correlation between the microscopic X-strain ε_1_ of the NMBS with the computational tensile X-strain ε_1_ (mean ± S.D.). **p.** Additionally, there is a strong correlation between the microscopic Y-strain ε_2_ of the NMBS with the computational compressive Y-strain ε_2_ (mean ± S.D.). **q.** Quantitative analysis of ROI 2 (region adjacent to the circular defect and parallel to the uniaxial tension) reveals a strong correlation between the microscopic X-strain ε_1_ of the NMBS with the computational tensile X-strain ε_1_ (mean ± S.D.). **r.** Additionally, there is a strong correlation between the microscopic Y-strain ε_2_ of the NMBS with the computational compressive Y-strain ε_2_ (mean ± S.D.).

To validate the accuracy of the NMBS strain mapping during uniaxial tensile testing, we correlated the average tensile and compressive strain observed via fluorescence microscopy to the measured macroscopic distances between fiducial marks on the surface of the PDMS test strip. We also measured the overall width and thickness of the test strip throughout tensile testing. The mean and standard deviation of the microscopic tensile ε_1_ (parallel to uniaxial testing) and compressive ε_2_ (perpendicular to uniaxial testing) strains of NMBS were plotted against the macroscopic tensile ε_1_ (Figure 2g) and compressive ε_2_ (Figure 2h) strain calculated from the fiducial marks and strip dimensions. In each case, the microscopic strain measurement was highly correlated with the macroscopic strain measurement (Pearson correlation coefficient of 0.999), establishing that the NMBS accurately tracks strain. Further, for the PDMS we measured the macroscopic compressive stretch ratios λ_2_ and λ_3_ and calculated the product of the macroscopic stretch ratios λ_1_λ_2_λ_3_. The stretch ratios λ_2_ and λ_3_ were highly correlated (Pearson correlation coefficient of 0.998, slope = 1) (Figure 2i), and the product of the macroscopic stretch ratios λ_1_λ_2_λ_3_ was equal to 1 at all steps of the uniaxial stretching (Figure 2j), indicating that PDMS behaves as an isotropic incompressible material in agreement with the literature.^34^ To evaluate the reproducibility we performed uniaxial tensile testing with 45% strain loading-unloading cycles at 3% increments, the microscopic tensile ε_1_ and compressive ε_2_ strain measurements remained highly correlated with the macroscopic tensile ε_1_ and compressive ε_2_ strain measurements (Pearson correlation coefficient of 0.999) (Figure S2).

For large uniform deformation macroscopic fiducial marks are sufficient for tracking strain during uniaxial tensile testing, but they do not provide the ability to measure heterogeneous strains at high resolution. To evaluate the NMBS for this purpose, we tracked microscopic strain fields surrounding a circular defect within a PDMS test strip during tensile testing and focused on two regions of interest (ROI 1, ROI 2) (Figure 2k). Following 40% uniaxial strain, ROI 1 lateral to the defect showed X-axis tensile strain ε_1_ was increased next to the defect, and Y-axis compressive strain ε_2_ showed the same behavior (Figure 2l). ROI 2 longitudinal to the defect showed decreased X-axis tensile strain ε_1_ and Y-axis compressive strain ε_2_ next to the defect (Figure 2m). To validate these experimental results using the NMBS at the microscale, we used FEA to model uniaxial testing of the PDMS with a 0.4 mm circular defect, taking advantage of the PDMS having well characterized mechanical properties and behaving as an ideal elastomer. Simulations of the uniaxial tensile testing confirmed our experimental results showing increased X-axis tensile strain ε_1_ and Y-axis compressive strain ε_2_ near the defect in ROI 1, and decreased X-axis tensile strain ε_1_ and Y-axis compressive strain ε_2_ near the defect in ROI 2 (Figure 2n). Importantly, there was high correlation (Pearson correlation coefficient of 0.999) between the microscopic observed using the NMBS and FEA-simulated strain within the X-axis and Y-axis for both ROI 1 and ROI 2 (Figure 2o, p, q, and r). Taken together, these data demonstrate the ability of the NMBS to accurately track and quantify homogenous and heterogeneous strains at the microscale.

As a final validation step, we wanted to use the NMBS to track strain on a biologically derived compressible hydrogel and engineered a fibrin tensile test strip for this purpose. Fluorescence images of the NMBS in its reference state (Figure S3a) and following 50% uniaxial strain (Figure S3b) demonstrate the ability of the NMBS to adhere to and track microscopic strain within a fibrin hydrogel. Quantification of individual segment strain and contour mapping reveal an average tensile strain of 0.4938 ± 0.0238 and an average compressive strain of 0.5188 ± 0.0403 following 50% uniaxial strain of the fibrin test strip (Figure S3c and d). In agreement with our results for PDMS test strips, for fibrin-based test strips, the microscopic strain measurement via the NMBS was highly correlated (Pearson correlation coefficient of 0.999) with the macroscopic strain measurements further confirming that the NMBS accurately tracks mechanical strain (Figure S3e and f). Unlike PDMS, fibrin test strips stretch ratios λ_2_ and λ_3_ were not equivalent (Figure S3g; slope = 1.42), and the product of the macroscopic stretch ratios λ_1_λ_2_λ_3_ was less than 1 at all steps of the uniaxial stretching (Figure 2h) indicating that fibrin behaves as a compressible material^35^. Taking the fibrin and PDMS data, these data show that the NMBS can accurately track strain and characterize different material properties.

### Mapping Subcellular, Cellular and Multicellular Strain with the Nanomechanical Biosensor

To apply the NMBS to biological systems we started at the single cell level. Traction force measurements have become a widely used methodology, but measurements are largely limited to forces applied to the cell-substrate interface.^26,27, 36–38^ The NMBS can be applied to the apical surface of the cell to provide new information on membrane deformation. To demonstrate this, the NMBS was applied to the apical surface of human skeletal muscle cells (HSMC) grown in culture using the 2 µm x 2 µm mesh to provide subcellular resolution. HSMCs were labeled with CellTracker™ Green for visualization (Figure 3a) and the fluorescently-labeled NMBS conformed to and adhered to the cell surface (Figure 3b). A custom image analysis pipeline was created using Imaris image analysis software and MATLAB software to extract the 3D strain information from the NMBS deformation over time (Figure S4). As a representative HSMC migrates over a 20 min period, tensile strain increases at the trailing edge and compressive strain increases at the leading edge (Figure 3c). Tracking of and visualization of displacement vectors for each node within the NMBS reveals the magnitude and direction of movement for the apical membrane surface in 3D (Figure 3d). Further analysis of the region of tensile strain at the trailing edge (ROI 1) and the region of compressive strain at the leading edge (ROI 2) revealed average peak tensile strains of 110% ± 22% and compressive strains of 39% ± 9% (Figure S4). Utilizing the NMBS to track cellular strain thus allows for the creation of an apical surface strain map with subcellular resolution.

**Figure 3.**
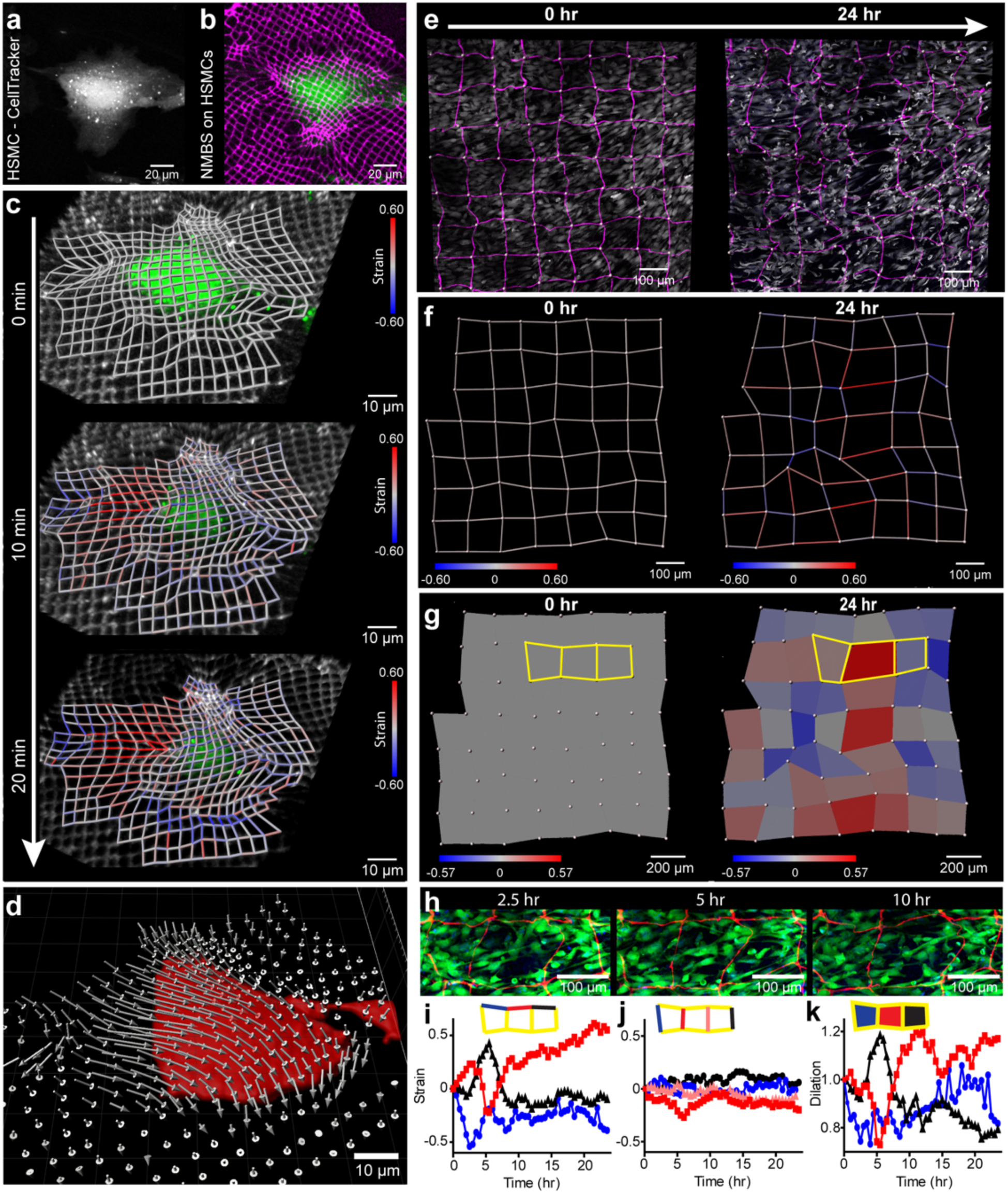
3D cellular strain mapping with the NMBS. **a.** Human skeletal muscle cells (HSMC) visualized with CellTracker-488. **b.** Alexa-647 fibronectin NMBS (2 µm X 2 µm X 2 µm, magenta) applied to the HSMC (green). **c.** Mapping of the NMBS strain over time reveals local regions of tension and compression following cell migration and contraction. **d.** 3D displacement vector map (gray arrows) showing the magnitude and direction of displacement for all NMBS mesh nodes as the cell compresses (Red surface) over 20 min. **e.** Alexa-555 fibronectin NMBS (100 µm X 100 µm X 10 µm, magenta) applied to C2C12 cells visualized with CellTracker-488 (gray). Images were acquired every 30 min for 24 hours to track strain following transition to C2C12 differentiation media. **f.** Strain mapping of the NMBS (red = tension, blue = compression) following transition to differentiation media reveals localized regions of tension and compression. **g.** Area dilation was calculated for each section of the NMBS over time and shows spatial regions of expansion and contraction. **h.** Regions of interest extracted showing cellular elongation events at 2.5, 5, and 10 hours following transition to differentiation media (NMBS, red; C2C12 cells, green). **i, j.** Quantification of segments from the region of interest in “g” during the C2C12 differentiation process shows region specific strain rates during the 24-hour time course, with the largest magnitude strains occurring in the x-direction. **k.** Time-dependent mapping of regional dilation during the C2C12 differentiation process for ROI analyzed in i and j.

To expand the technology to multicellular measurements we used the NMBS to track strain in C2C12 mouse myoblast monolayers, with the goal to study myoblast migration and fusion of single cells into multinucleated myotubes. The NMBS was engineered as a mesh with 10 µm wide fibers spaced at 100 µm, applied it to a confluent C2C12 monolayer, and the media was changed to differentiation media. Following 24 hours of live imaging in differentiation media, the C2C12 cells labeled with CellTracker™ Green were observed to undergo coordinated and regionally localized migration into areas of high and low cell density, highlighted by the deformations within the NMBS (Figure 3e). Strain between nodes of the mesh quantified and mapped from the NMBS deformations showed large tensile and compressive strains correlating with the regions of low and high cell density, respectively (Figure 3f). Analysis of dilation for the area within each square of the NMBS mesh revealed a field map of the expansion and contraction (Figure 3g). To look at the correlation between strain and the dynamics of cell migration and elongation, we further evaluated a region of interest that exhibited high tensile strain and large dilation following 24 hours, which showed oscillations over time (Figure 3h). Quantification of NMBS segments within this region revealed segment-specific oscillating changes in the X-strain (Figure 3i), with minimal change in the Y-strain (Figure 3j). Dilation analysis of this region was correlated with the X-strain data (Pearson Correlation 0.806 ± 0.094), but not the Y-strain data (Pearson Correlation −0.206 ± 0.231) suggesting that the regional expansion and contraction observed during the C2C12 migration and elongation is driven by uniaxial X-axis biased tension and compression (Figure 3k).

### Dynamic Strain Measurement During Cardiomyocyte Contractions

In many biological systems, cells and tissues undergo dynamic movements that drive developmental and physiological processes. This is especially true for cardiac tissue, where cardiomyocytes (CMs) display coordinated contraction and electromechanical coupling into a syncytium.^39^ For human CMs derived from embryonic stem cells (hESC), coordinated contraction and shortening are important markers of maturation and disease phenotype^40,41^ and heterogeneity following differentiation is also a critical quality metric but difficult to measure in 2D culture settings.^41^ Here we assessed hESC-CM function using the NMBS to measure CM beat frequency and contraction induced strain, combined with fluorescent imaging of calcium signaling. We used an NMBS mesh with 10 µm wide fibers spaced at 100 µm applied to a confluent hESC-CM monolayer incubated with Fluo-4 calcium indicator dye (Figure 4a). Motion of the NMBS was used to extract CM beat frequency from each pixel region of the NMBS by measuring changes in average fluorescence intensity over time (Figure 4a and b). A fast Fourier transform (FFT) analysis was performed to extract the dominant frequency from the fluctuations in average fluorescence intensity over time (Figure 4c). Using these values an image can be reconstructed as a region-specific CM beat frequency mapped to each pixel of the NMBS in the field of view (Figure 4d). To validate this approach, the beat frequency of spontaneously contracting CMs calculated from NMBS motion was compared to the calcium signaling and found to be the same (Figure S5a). Similar results were found when paced with either 1 Hz or 2 Hz field stimulation the NMBS and calcium wave frequency data was identical and perfectly matched the stimulation rate (Figure S5b). Mapping the beat frequency on to the NMBS showed that it was uniform across the field of view at 2 Hz stimulation (Figure S5c).

**Figure 4.**
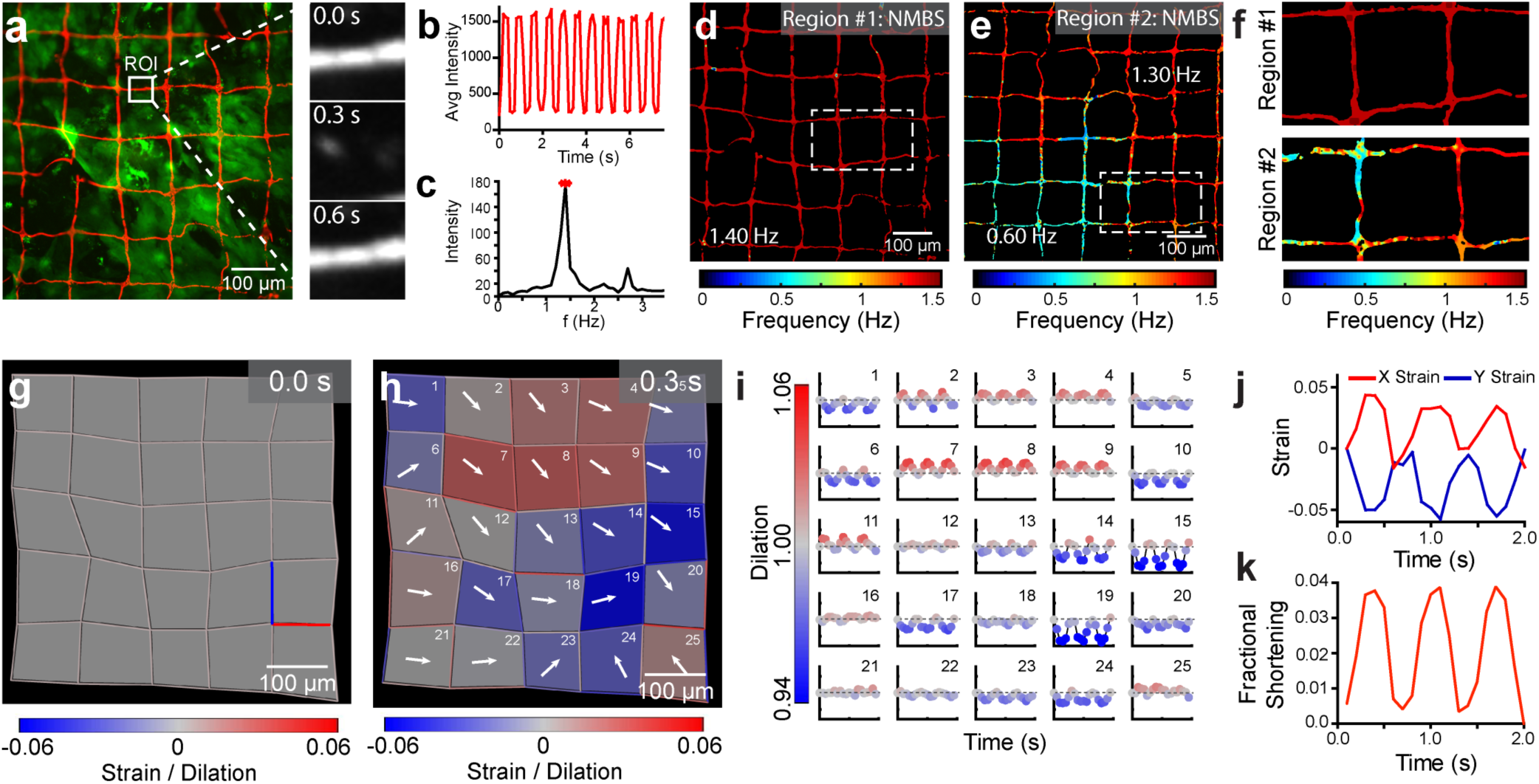
Beat frequency analysis and strain mapping during cardiomyocyte contractions. **a**. Fluorescence image of representative Alexa-555 fibronectin NMBS (Red, 100 µm X 100 µm X 10 µm) applied to iPSC-derived cardiomyocyte monolayer loaded with Fluo-4 calcium indicator dye (Green). White box indicates ROI for NMBS motion analysis. Motion analysis of ROI displays NMBS segment moving in and out of the ROI over time. **b.** Quantification of the average fluorescence intensity from the ROI through time. **c.** Example power spectral density plot from the Fourier transformed data of a NMBS ROI segment showing the intensity at each frequency with a dominant frequency of 1.4 Hz. **d.** A synthetic image is constructed containing the frequency information of every pixel within the NMBS to visualize variability of contractile frequencies for cardiomyocyte monolayers. A dominant uniform frequency of 1.4 Hz is overserved. **e.** Example region #2 of contractile cardiomyocytes demonstrating heterogeneous beat frequencies of 1.3 and 0.6 Hz. **f.** Magnified images of region #1 and #2 demonstrating differences in heterogeneity between fields. **g,h.** Time-series showing regional dilation and NMBS segment strain throughout a contraction cycle of diastole and systole. Displacement vectors (white arrows) for each square region defined by the NMBS show the contractile direction between peak systole and diastole. **i.** Analysis of dilation over a 2 second time interval defined by the NMBS meshed regions (Red = tension, Blue = compression). **j.** Quantitative analysis of example X (red line in panel g) and Y (blue line in panel g) strain from NMBS segments shows the tensile and compressive strain rate over time for cardiomyocytes with a beat frequency of 1.4 Hz, maximum tension of 4% and compression of 6%. **k.** Fractional shortening is calculated from a regional dilation analysis of ROIs 13,14,18, and 19 demonstrating a peak fractional shortening of 3.8% ± 0.06 between diastole and systole (mean ± S.D.; n=3).

Next, hESC-CM cultures were evaluated for beat frequency heterogeneity using the NMBS mapping. In many instances, the typical field of view revealed homogeneous contractions (Figure 4d), which is indicative of good cell-cell coupling between CMs and the formation of a continuous syncytium. However, there were also fields of view that had heterogeneity (Figure 4e) with variable beat frequencies ranging from approximately 0.5 Hz to t1.5 Hz. Transitions between beat frequency rates appeared to occur at sharp boarders within continuous cell monolayers and were seen to differ between fields of view within a single coverslip (Figure 4f). To obtain enhanced resolution for beat frequency analysis, we also used an NMBS mesh with a smaller gird size consisting of 10 µm wide fibers separated by 20 µm and compared this to the calcium signaling. The increased resolution provided by the denser NMBS mesh pattern together with the calcium signaling revealed that the heterogeneity, in terms of large differences in beat frequency, were due to other cell types that were not hESC-CMs in the culture. Specifically, regions that had beat frequencies from 0 to 5 Hz did not show calcium signaling (Figure S5d), indicating that these were not hESC-CMs. To support this, field of views that had uniform beat frequency determined by the NMBS also showed uniform calcium signaling. Differentiation of hESC-CMs is ~90% efficient so non-contractile cells are expected. The advantage of the NMBS over calcium imaging, however, is that we can assess beat frequency and heterogeneity without needing to use an intracellular fluorescent stain that has the potential to cause phototoxicity.

In addition to beat frequency, the NMBS can be used to measure strain and regional dilation between diastole (full relaxation) and systole (peak contraction). We observe that even in a region with uniform beat frequency, the monolayer of hESC-CMs show a large degree of variability in the direction of motion and magnitude of expansion or contraction (Figure 4g and h). Plotting the dilation for each grid as a function time (Figure 4i) confirms that all areas are contracting in synchrony. We observed peaks in expansion (Region #8) of 1.04 and contraction (Region #19) of 0.931 (Figure 4i), while other areas had relatively small changes (Region #21). It appears that the clusters of hESC-CMs that cause these different dilation patterns are larger than the grid size, as the expansion or contraction typically involves multiple adjacent regions. Further, segments connecting the nodes of the NMBS mesh can be used to determine the strain in the X and Y direction at different regions of the culture (Figure 4j). To determine fractional shortening during hESC-CM contraction, we calculated the percent change in area between diastole and systole for a combined region (Region #13,14,18, and19). The mean fractional shortening following three contractile cycles was 3.84% ± 0.06% (Figure 4k), this measurement is the deformation of the apical surface of the hESC-CMs, as the basal surface is adhered to a coverslip. These data demonstrate the versatility of the NMBS for quantifying heterogeneity in the contractility of hESC-CMs that would not be evident from voltage or calcium imaging.

### Mapping 3D Strain in Developing Tissue

Developing tissue undergoes dynamic changes during morphogenesis as a result of cell-generated mechanical forces. Studying these changes in a direct manner is difficult without interfering with the internal structure and function of the tissue. The NMBS offers a solution to this problem through application of a nanometers thick FN mesh onto the surface. To demonstrate this we used the Drosophila ovariole as a developmental model to investigate two aspects of mechanical strain; slow dynamics associated with cytoskeletal rearrangements, and fast oscillations associated with muscular contractions^42^. Ovarioles expressing moesin-GFP to visualize the actin cytoskeleton and histone-2B-RFP to visualize the nucleus were dissected for germarium through stage 6 ovariole chambers,^43^ transferred to a glass coverslip, and an NMBS with 2 µm wide lines at 2 µm spacing was applied (Figure 5a). The NMBS adhered to the 3D surface topology of the ovarioles and appeared to make conformal contact throughout, based on cross-sections from confocal images (Figure 5b). A 4-hour time-lapse of a stage 4 ovariole (region indicated by dashed white box in Figure 5a) showed changes in strain magnitude and direction over time. At both 20 minutes and 2 hours and 40 minutes into the imaging series, a large band of tensile strain appeared along the top edge of the stage 4 ovariole (Figure 5c). Closer investigation into this region (dashed white box in Figure 5c) showed an increase in moesin-GFP intensity associated with the increase in tensile strain (Figure 5d). The average strain for multiple segments in this region (white arrow heads in Figure 5d) and moesin-GFP fluorescence intensity plotted as a function of time revealed a peak in moesin-GFP fluorescence that corresponded to a peak in tensile strain that steadily decreased until the next peak in moesin-GFP fluorescence (Figure 5e).

**Figure 5.**
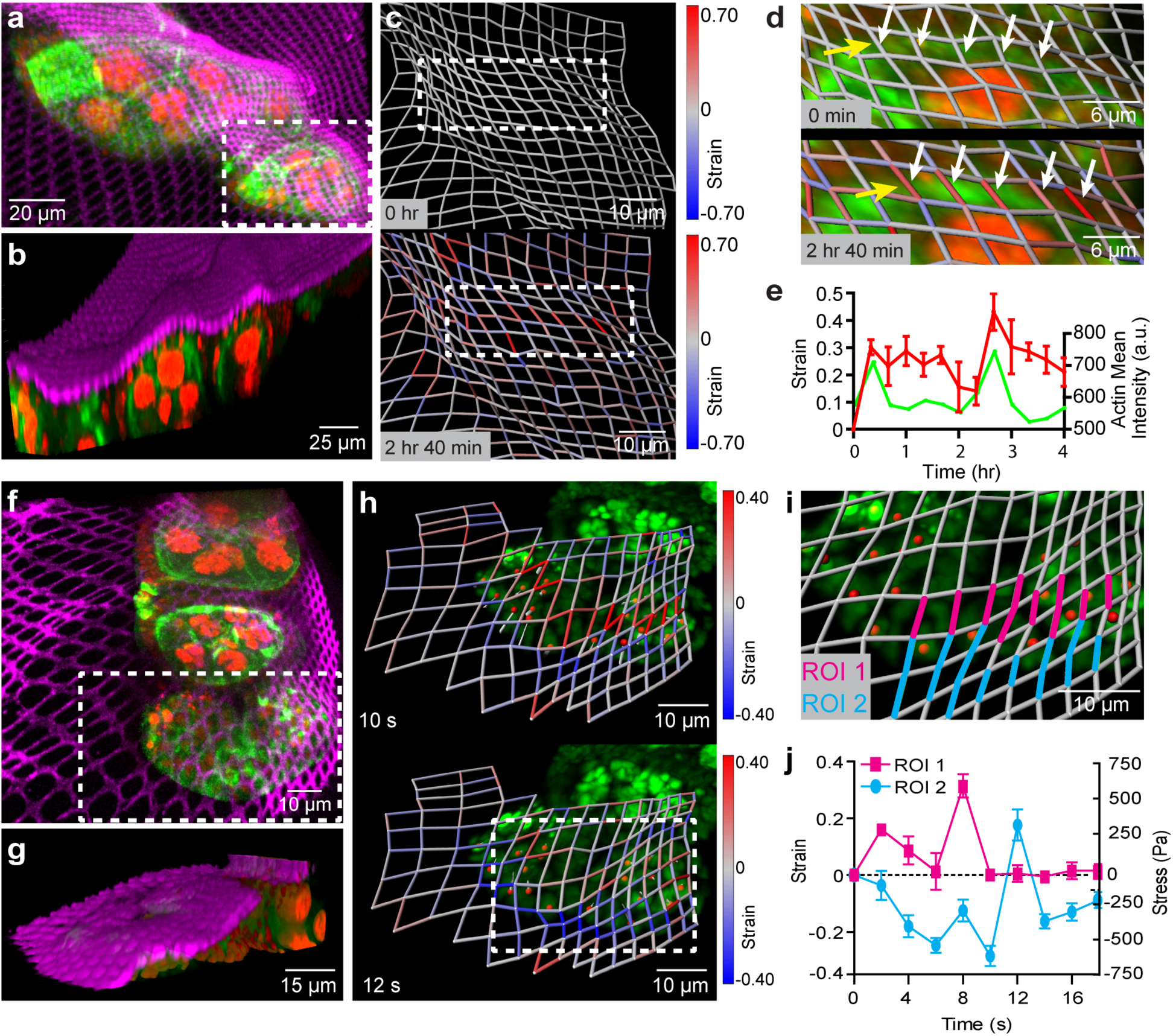
Utilizing the NMBS for 3D strain mapping on drosophila ovariole tissue. **a.** Maximum intensity projection of fluorescence image from live confocal imaging of Alexa-647 fibronectin NMBS (Magenta, 2 µm X 2 µm X 2 µm) applied to a developing drosophila ovariole expressing moesin-GFP (Green) and Histone-2B-RFP (Red). **b.** 3D render of a diagonal cut plane demonstrating the ability of the NMBS to conform to the 3D surface of the ovariole. **c.** Strain analysis and quantification of region in panel a (white box) at two time points show regionally localized tensile (red) and compressive (blue) strain over time. **d.** Magnified images (white box in panel c) of high tensile region (white arrows) corresponding to localized moesin-GFP fluorescence (green, yellow arrows). **e.** Quantification of the average tensile strain (red, white arrows) within the magnified region shows strain peaks correlate with increased moesin-GFP fluorescence (green, yellow arrows) over a 4-hour time course (mean ± S.D.; n = 5). **f.** Maximum intensity projection of fluorescence image from live high-speed spinning-disk confocal imaging of Alexa-647 fibronectin NMBS (Magenta, 2 µm X 2 µm X 2 µm) applied to a contracting drosophila ovariole expressing moesin-GFP (Green) and Histone-2B-RFP (Red). **g.** 3D render of a diagonal cut plane demonstrating the ability of the NMBS to conform to the 3D surface of the ovariole. **h.** Time-lapse imaging of region in panel f (white box) at 0.5 Hz of a 20 µm section of a contracting ovariole with the NMBS (magenta) and histone-2B-RFP (green). Regions of tension (t = 10s, max of 1.55) and compression (t = 12s, max of 0.63) are observed during contraction and extension respectively. As the nuclei translate upward during ovariole contraction (t = 10 s; red dots, gray commit tails) there is an increase in tension at their leading edge (max tension = 1.44). As the nuclei move down and the ovariole compresses (t = 12 s), a band of compression along the bottom edge of the ovariole is observed (max compression = 0.60). **i.** Magnified image of strain analysis showing the 3D deformation during ovariole contractions and highlighting regions of compression and tension for quantification. **j.** Quantification of the maximum tensile and compressive regions within the strain map, (ROI 1, magenta; ROI 2, cyan) of the contracting ovariole. Stress during contraction was calculated by using values for the elastic modulus of the germarium and stage 1 ovariole of 2 kPa (mean ± S.D.).

We also explored the ability for the NMBS to track fast dynamic strains over short time scales using high-speed spinning disk 3D confocal imaging (Figure 5f). Our goal was to measure the magnitude of tensile and compressive strain during contractions and correlate the peaks in strain with the magnitude and direction of nuclear motion. Analysis of the NMBS region covering the germarium through stage 4 ovariole revealed distinct patterns of tensile (Figure 5h, top) and compressive (Figure 5h, bottom) strain. These peaks in strain were highly correlated with the coordinated nuclear movement accompanying the muscle sheath contraction (Figure S6). Quantification of the regions comprising peak tensile (ROI 1) and compressive strain (ROI 2) demonstrated maximum tensile strain of 32.88 ± 8.53% and a maximum compressive strain of 26.73 ± 9.14% (Figure 5j) during a contraction. Additionally, by estimating the elastic modulus of the stage 4 ovariole as 2 kPa ^44^, we calculated the stress the muscle contractions exert on the ovariole based upon the NMBS deformation. We observed a peak tensile stress of 657.6 ± 170.6 Pa and peak compressive stress of 534.6 ± 182.8 Pa. Additional experiments looking at the frequency of ovariole muscular contractions confirmed a contraction frequency of 1 per 12 seconds and exhibited similar maximum tensile and compressive strain during contractions of 42.95 ± 9.43% and 22.63 ± 8.19% respectively (Figure S6).

## Discussion

Our results demonstrate that an ultra-thin, fluorescently-labeled FN mesh of tunable dimensions can serve as a NMBS to accurately quantify and map strain from subcellular to multicellular length scales. The NMBS is unique in that it can be directly applied to the surface of a range of biological and non-biological materials acting as a fiducial marker for strain analysis by quantifying deformation over time. Additionally, by choosing the appropriate fluorophore to conjugate to the NMBS, it can be combined with 3D confocal or multiphoton imaging allowing for simultaneous monitoring of biological processes and strain measurements.

The NMBS is unique in that the resolution and geometric pattern can be tuned to fit a range of experimental needs. Our original design consisted of a square lattice mesh 1 cm^2^ in overall area, providing adequate coverage for many biological samples. However, this can be scaled up or down to suit a particular sample and other patterns (e.g., circular holes, hexagonal arrays, triangular mesh, auxetic lattices) can be used to optimize or enhance fabrication, detection, and analysis in a tissue specific manor^31^. In this manuscript we choose to focus on the square lattice template as a proof of principle and will investigate additional NMBS designs in future work. Our results reveal that a NMBS with mesh lattice a 2 µm line width and 2 µm spacing enables subcellular measurement of strain. At this size, we maintain robust fabrication of the NMBS while also providing a strain map with single micron scale resolution. It is possible to create an NMBS at resolution below 2 µm; however, utilization of chrome photomasks and nanofabrication facilities are required as the pattern size decreases. Mesh lattices with 10 µm wide lines and 100 µm spacing was found to be a good compromise between resolution and coverage area for multicellular tracking of myoblast differentiation and cardiomyocyte contractility.

Additionally, by creating a strain sensor entirely comprised of ECM proteins such as FN, there is no exogenous material restricting cell growth or migration. For cells and tissues that do not express appropriate integrins for FN binding, alternative ECM proteins can be conjugated with a fluorescent dye and used for NMBS fabrication. Aside from the ability to construct the FN NMBS, the SIA process allows for microcontact printing of nanometer thick collagen I, IV, and laminin^31,45^. However FN is an ideal ECM protein for NMBS fabrication because it is hyperelastic and can achieve strains over 700% making it appropriate for fiducial tracking of large strains associated with tissue formation^46^. By passively stretching with the underlying cells or tissue, the FN NMBS allows minimal disruption of developmental processes and native biomechanics. Additionally, as the FN NMBS is extremely thin at ~4 nm, it is unlikely to mechanically influence cells adhered to a rigid substrate or 3D multicellular tissues where there is typically a collagen containing ECM matrix that is significantly stiffer and more extensive than the NMBS mesh^47^. Thus, as the cells in 2D and 3D migrate, differentiate, and contract, the FN NMBS is easily deformable allowing visualization and calculation of dynamic 3D surface strain.

In developmental biology, measurement and identification of mechanical forces required for tissue formation are often limited to 2D indirect approaches and invasive or destructive techniques^23,25^. Recent advances in high speed confocal fluorescence imaging techniques have facilitated the development of new 3D biomechanical force sensors such as 3D traction force microscopy and functionalized oil microdroplets^21,27^. 3D traction force microscopy is an elegant solution for directly measuring subcellular strain within a substrate of known material properties, but analysis is computationally demanding and is difficult to extend beyond the cellular length scale. For functionalized oil microdroplets the sensor resolution depends on the size and deformability of the droplet to provide direct quantification of the inter-cellular mechanical force within a tissue, but sparse labeling is needed to avoid developmental tissue defects. The ability for the NMBS to directly measure surface strain over a spatial range from approximately 1μm to 1 mm in biologically relevant hydrogels and samples serves to bridge the gap and compliment these modalities. Application onto isolated drosophila ovarioles highlight the ability of the NMBS to track 3D tissue strain over time with subcellular resolution during development as well as throughout fast oscillating muscle sheath contractions. One can imagine that by combining fluorescence imaging-based approaches such as 2D traction force microscopy (cell-substrate interface), functionalized oil microdroplets (intercellular), and the NMBS (free apical surface), we could simultaneously map 3D biomechanical strain to better understand how force is transmitted within cells and tissue.

Currently, the NMBS platform maps strain by determining a change in length or area between defined geometric regions. In addition to mapping and quantifying strain over time, the NMBS can also approximate stress from strain if the samples elastic modulus is obtained via AFM, functionalized oil microdroplets, or other mechanical testing. This is a highly informative approach; however, the surface of a cell or tissue is a biomechanical continuum with regions undergoing dynamic non-uniform expansion and contraction. The use of the NMBS allows for a discretization of these continuous surfaces, sampling the surface at many points. If a dense enough mesh was used, the complete continuum of the surface of the cell could be estimated using interpolation between adjacent nodes in the NMBS^48^. This would allow for not only a complete strain analysis of the surface in 3D but also a measurement of the energy in the surface^49^. Combined with a constitutive law for the cellular or tissue surface, changes in strain energy could be derived from this continuum approximation^50^. Together, this highlights area of future extension of the NMBS technology and would allow for a more complete understanding of the stresses driving developmental morphogenesis or disease progression.

## Methods

### Fabrication of the nanomechanical biosensors

The NMBS were fabricated using an adaptation of previously reported surface-initiated assembly technique^31,51^. Briefly, square-lattice meshes (100 μm length X 100 μm width X 10 μm line thickness per grid segment) were first designed using AutoCAD software. The CAD file was then transferred to a transparency photomask (CAD/Art Services, Inc., Bandon, OR, USA), where the spaces and the segments of the square-lattice meshes were dark and transparent, respectively. Square glass wafers (Fisher No. 12-543-F 45×50×2) were spin-coated with Photoresist SPR-220.3 at 5000 rpm for 20 sec, then baked on a 115°C hot plate for 90 sec, then exposed to ultraviolet (UV) light through the transparency photomask, then baked on a 115°C hot plate for 90 sec, and finally developed for 1 min using MF-26A developer. Sylgard 184 (Dow Corning) polydimethylsiloxane (PDMS) elastomer was prepared per manufacturer’s directions by mixing 10 parts base to 1 part curing agent (by weight) using Thinky-Conditioning mixer (Phoenix Equipment Inc., Rochester, NY, USA) for 2 min at 2000 rpm followed by 2 min of defoaming at 2000 rpm. The PDMS was then cast over the topographically-patterned photoresist-coated glass wafer inside a petri dish and placed in a 65°C oven to cure the PDMS. Cuboid PDMS stamps approximately 1×1 cm were cut out of the ~5 mm thick PDMS layer.

To begin the surface-initiated assembly process, the PDMS stamps were sonicated in 50% ethanol for 30min, dried with nitrogen gas, and coated with a 50 μg/mL human FN solution (Corning). The FN solution contained a 40% (v/v) Alexa Fluor® 555, or 633 (ThermoFisher) C5 fluorescently labeled FN. After 1hr of incubation at room temperature, the FN-coated PDMS stamps were rinsed in sterile ddH2O, dried with nitrogen, and stamped onto 25 circular glass coverslips (Fisher No. 12-545-86 25CIR-1D). Prior to stamping, the coverslips were cleaned and spin-coated at 6000 rpm for 1 min and 37 sec with 2% g/mL poly(N-isopropyl acrylamide) (PIPAAm) (Scientific Polymer, mw 300,000) diluted in 1-Butanol. After 1 hour the PDMS stamps were carefully removed from the glass coverslips, leaving behind a microcontact-printed fluorescently labeled FN-based 2D square-lattice meshes or NMBS onto the sacrificial PIPAAm surface prior to release and subsequent transfer to different materials. Design and fabrication of the 20 μm length X 20 μm width X 10 μm line thickness NMBS and 2 μm length X 2 μm width X 2 μm line thickness NMBS was identical to the above-mentioned procedure.

### Creation of hydrogel test strips for uniaxial testing

For uniaxial testing, NMBS were integrated onto dog bone shaped tensile testing strips made of fibrin and PDMS elastomer and subjected to uniaxial mechanical strain. See Supplementary Figure 1 for dimensions of dog bone test strips which have been slightly adjusted from ASTM E8 to fit our experimental setup.

#### Fibrin Test Strips

For fibrin test strip fabrication, 40 mg/mL bovine fibrinogen (MP Biomedicals) was combined with 10 U/mL bovine thrombin (MP Biomedicals) at a final concentration of 36 mg/mL and 1 U/mL, respectively, to catalyze the conversion of fibrinogen to fibrin^52^. The fibrin mixture was casted into 3D printed plastic dog bone molds containing small Velcro pieces (VELCRO® Super-Grip Double-Head Hook) inserted at both ends to grip the gel during tensile testing. Molds were sprayed with Teflon (WD-40® SpecialistTM Dirt & Dust Resistant Dry Lube Spray) to facilitate mold release following casting. After 30 minutes at room temperature, the fibrin hydrogel solidified.

#### PDMS Test Strips

For PDMS test strip fabrication, Sylgard 184 and Sylgard 527 (Dow Corning) were blended at a mass ratio of 5:1 as previously described by our group^45^. Briefly, Sylgard 184 and 527 were prepared per manufacturer’s directions and mixed in a Thinky-Conditioning mixer for 2 min at 2000 rpm followed by 2 min of defoaming at 2000 rpm. Once mixed, the 5:1 PDMS 184:527 was casted into 3D printed dog bone molds as previously described. The molds were placed in a 65°C oven overnight to ensure proper PDMS curing. Prior to NMBS application, the PDMS test strips were treated with 15 minutes of UV Ozone to create a hydrophilic surface to facilitate NMBS attachment.

### Uniaxial tensile testing

To transfer the NMBS to the hydrogel test strips, the hydrogel test strip was pressed onto the PIPAAm-coated glass coverslip stamped with the fluorescent NMBS. Warm (37°C) 1X PBS was added to the glass coverslip and test strip. As the temperature dropped below the lower critical solution temperature of PIPAAm (~32°C), the gradual dissolution of PIPAAm resulted in release of the coverslip and integration of the NMBS onto the bottom surface of the hydrogel test strip.

Fibrin and PDMS test strips containing an applied Alexa-555-FN-NMBS were mounted to micromanipulators (Eppendorf TransferMan NK) are both ends using fast hardening two-part epoxy (Devcon high strength 5min epoxy) through the embedded Velcro attachment points. Uniaxial testing was performed on top of a Nikon Eclipse TI wide-field fluorescence microscope. Fluorescence imaging was performed with a CoolSnapES (Photometrics) camera, X-Cite 120PC light source (Excelitas Inc.), and appropriate excitation and emission filters for Tx-Red fluorescence. Fiducial marks spaced 8 mm apart were introduced in the gauge region of the test strip on the top surface to provide tracking of macroscopic deformation. Uniaxial tensile testing was performed in 100 μm increments. Fluorescence images of the NMBS (10X air objective) and macroscopic images (GoPro HERO3 camera) of fiducial marks were acquired at each step. Macroimaging of the test strip side edges provided tracking of compressive strain ε_2_ at the macroscopic scale. Differential interference contrast (DIC) imaging of the test strip top and bottom surface provided thickness measurements during mechanical testing for calculation of macroscopic compressive strain ε3 along the z-axis.

For uniaxial testing involving the circular defect, a circular defect of diameter 0.4 mm was introduced in the center of a 555-FN-NMBS PDMS test strip gauge region using a hole punch tool. Test strips with circular defects were mounted onto micromanipulators for uniaxial testing as previously described.

### 2D Image analysis for quantification of NMBS strains

MATLAB-based analysis code was written to measure the changes in NMBS segment length from the imaging data acquired during uniaxial tensile tests to calculate the NMBS microscopic 2D strains. Figure 2d shows a representative microscopy image of NMBS at macroscopic tensile strain of PDMS equal to 0.45. First, the image series was imported into MATLAB, background subtracted, and thresholded to convert the NMBS fluorescence into a binary image. The binary image was skeletonized to identify the intersection nodes. Following node-to-node pair assignment lengths between each node pair was calculated and overlaid onto the original NMBS image (Figure 2e). To quantify strain during uniaxial testing, the segment lengths between node pairs were calculated for each testing interval and converted into engineering strains 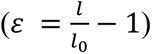 with respect to their undeformed reference length. Positive values of strain corresponded to tensile components of strain, while negative values of strain corresponded to compressive components of strain (Figure 2f). To visualize the strain fields, we converted the strain values fxgor each segment into a color-coded map for both tensile and compressive mechanical strains of NMBS (Figure 2f).

### ANSYS Mechanical Simulations

To validate the strain results obtained by the NMBS during uniaxial tensile testing we performed finite element analysis to simulate the tensile and compressive strains for a PDMS strip with a 0.4 mm circular defect. Fusion 360 modeling software was employed to create an *in-silico* representation of the tested PDMS strip with the circular defect. Of note, the model was made longer than the sample tested on the stretcher. This was done to negate the edge effects of having a fixed boundary condition. Once modeled, the object was exported as an .STL and imported into the 2019 ANSYS Mechanical static structural simulation.

A Mooney-Rivlin 2 parameter model was assigned to the object. The general form of this model is *W* = *C*_1_(*I*_1_ − 3) + *C*_2_(*I*_2_ − 3) where W is the energy strain function, I1 and I2 are the first and second invariants of the LaGrange-Green deformation tensor. Furthermore, the material was assumed to be isotropic. *C*_1_ and *C*_2_ were assigned values of 0.0213 MPa and 0.0021 MPa, respectively for PDMS with a 5:1 mass ratio of Sylgard 184:527^45^. These parameters were determined by fitting experimental uniaxial stress-strain curves from a solid strip of our PDMS. An incompressibility parameter of 0.0001 *Mpa*^−1^ was assigned to enforce isochoric behavior of the model. A fixed boundary condition was applied to one face of the object. On the opposite face, a displacement boundary condition was set to produce a 60% strain of the entire sample, recapitulating the maximum strain applied during experimental uniaxial testing. After the simulation converged, global short axis (X) and long axis (Y) strains were calculated and rendered.

### Application of NMBS onto living cells and tissue

After 1 hour of microcontact printing the PDMS stamps were removed from the glass coverslips leaving behind the NMBS patterned onto the sacrificial PIPAAm surface. To apply to cells, tissue, or other substrates the NMBS is transferred to sterile gelatin carriers which melt upon incubation at 37°C to integrate the NMBS on the sample.

#### Preparation of sterile gelatin carriers and transfer of NMBS

Gelatin type A carriers (20% w/v) were prepared by casting into a circular silicon mold. The silicon molds were removed, and the gelatin gels were UV treated for 10 min to sterilize. NMBS-patterned PIPAAm-coated glass coverslips were placed with NMBS facing down onto the gelatin carriers inside the cell-culture hood for 1 minute. Sterile room temperature ddH2O was added between the glass coverslips and the gelatin carrier to cause dissolution of the PIPAAm and subsequent release and transfer of the NMBS to the top surface of the gelatin. Following transfer, aspirate all remaining water surrounding the NMBS-gelatin carrier and store in a sealed container at 4°C until ready for tissue application.

#### Application of the NMBS onto cells

Once transferred to the gelatin carrier, the NMBS-gelatin carrier is ready for application onto cells or tissue. For application onto cells, cells were grown on a glass coverslip and transferred to a metal imaging chamber (Attofluor Cell Chamber, ThermoFisher #A7816). Before the addition of media, the NMBS-gelatin carrier was lifted and applied to the cells with the NMBS side down. Light pressure was applied to the gelatin carrier to fully adhere the NMBS to the cells. Subsequent incubation at 37°C for 5 minutes melted gelatin and integrated the NMBS onto the top surface of the cells. Without aspirating the melted gelatin, fresh cell culture media was added and incubated at 37°C for 5 minutes. Prior to imaging, the media was changed to remove the melted gelatin and label cells with CellTracker™ Green CMFDA Dye (Invitrogen) and 2 μg/mL Hoechst 33342 dye (ThermoFisher Scientific) to stain for the cell nuclei. A final media change was performed to phenol red free CO2 independent Lebovitz L-15 (Invitrogen) media containing appropriate serum and nutrients for each cell line.

#### Application of the NMBS onto living tissue

For application to Drosophila ovariole tissue, the ovarioles were fanned out onto a glass coverslip and allowed to partially dry. The coverslips were then transferred to a metal imaging chamber (Attofluor Cell Chamber, ThermoFisher #A7816) and the NMBS-gelatin carrier was lifted and applied to the top side of the ovarioles with the NMBS side down. Light pressure was applied to the gelatin carrier to fully adhere the NMBS to the ovariole surface. Subsequent incubation at 37°C for 3 minutes followed by Hoechst 33342 containing Schneider’s media and incubation at 28°C for 10 minutes melted gelatin and integrated the NMBS onto the top surface of the ovarioles. A final media change was performed with fresh phenol-red free Schneider’s media before imaging at 25°C.

### Cell lines and cell culture

For our investigations of cellular strain using the NMBS we used human skeletal muscle cells (HSMC, Cook MyoSite), C2C12 mouse myoblast cells (CRL-1772, ATCC), and HES3 human embryonic stem cell (hESC)-derived cardiomyocytes. The HSMCs were grown in MyoTonic Growth Medium Kit (MK-4444, Cook MyoSite) with 1% (v/v) penicillin-streptomycin (15140-122, Life Technologies) at 37°C with 5% CO_2_ and maintained at ≤ 50% confluency following the manufacturers recommended protocol^53^. Prior to NMBS application and live fluorescence imaging, HSMCs were plated onto 25 mm glass coverslips and allowed to grow for 24 hours.

Culture and passage of C2C12 cells was performed in DMEM-high glucose (15-013-CM, Corning CellGro) supplemented with 10% (v/v) Fetal Bovine Serum (FBS) (89510-186, VWR), 1% (v/v) L-glutamine (25030-081, Life Technologies), and 1% (v/v) penicillin-streptomycin (15140-122, Life Technologies) at 37°C with 10% CO_2_ and maintained at ≤ 80% confluency. Prior to NMBS application and live fluorescence imaging, C2C12 cells were plated onto 25 mm glass coverslips, allowed to grow for 24 hours. To begin myoblast differentiation and myotube formation cells were transitioned to medium consisting of DMEM-high glucose supplemented with 2% (v/v) heat-inactivated horse serum (H1138 Sigma-Aldrich), 1% (v/v) L-glutamine (25030-081, Life Technologies), and 1% (v/v) penicillin-streptomycin (15140-122, Life Technologies).

Human cardiomyocytes derived from HES3 human embryonic stem cells were cultured and differentiated according to established and previously published protocols^54,55^. Briefly, hESCs were grown in Essential 8 (E8) medium (A1517001, Life Technologies) supplemented (S1459, Selleck Chemicals) in 6-well plates coated with Geltrex (12 µg/mL, A1413301, Life Technologies). Once 50% confluency was achieved, cardiomyocyte differentiation was initiated as previously^54,55^. Following differentiation and lactate-based metabolic purification^56^, spontaneously beating cardiomyocytes were maintained in CDM3 medium for up to 28 days prior to experimentation^57^. Prior to NMBS application and live fluorescence imaging, purified cardiomyocytes were plated onto 25 mm glass coverslips coated with Matrigel (12 µg/cm^2^, 356231, Corning) and maintained in CDM3 medium.

### Live Cell Fluorescence Microscopy

Various confocal fluorescence microscopes were utilized to acquire 3D time-lapse images of the NMBS applied to cells and tissue. For imaging of single cells (HSMCs, Figure 3a) and drosophila ovariole growth (Figure 5a) we performed imaging on an upright Nikon FN1 resonant scanning confocal microscope with a 40X CFI Apo NIR W objective, 4 laser lines (405, 488, 555, 633), 4 internal detectors (2 PMTs, 2 GaAsP), and a heated stage platform to maintain the temperature at 37°C. For drosophila ovarioles the temperature was maintained at 28°C. 3D image stacks were acquired for each time point in NIS Elements using a 1 μm z-step interval. For the HSMCs and drosophila ovarioles, 3D image stacks were acquired every 10 and 20 minutes respectively for the duration of the experiment.

To evaluate NMBS strain during the initiation of C2C12 cell differentiation we performed live confocal fluorescence imaging on an inverted Zeiss 700 LSM microscope equipped with a 20X Plan Apo objective, 4 laser lines (405, 488, 555, 633), 2 internal spectral PMT detectors, and a live cell incubation system to maintain the temperature at 37°C and CO2 at 5%. 3D image stacks were acquired for each time point in Zeiss Zen software using a 1 μm z-step interval. For the C2C12 cells, 3D image stacks were acquired every 30 minutes to limit the light exposure for the duration of the 24-hour experiment.

In order to visualize and track the 3D strain during drosophila ovariole muscle sheath contractions we utilized spinning disk confocal fluorescence microscopy. Specifically, imaging was performed using a XDi spinning disk confocal system (Andor) on a Nikon TiE-2000 inverted microscope with a 40X Apo LWD WI objective. The imaging system also consisted of fully temperature and humidity control to maintain 25°C, a mechanical Piezo XYZ-stage (Nikon), iXon 897 Ultra EMCCD camera (Andor), a 5 line laser combiner (405, 488, 515, 555, 647; Andor), a Lambda 10-3 filter wheel (Sutter), IQ2 imaging software (Andor), and an active air isolation table (TMC). High speed 3D image stacks were acquired for each time point in IQ2 software using a 0.2 μm z-step interval. For the fast contractions, 2 and 3 fluorescence channel 3D image stacks were acquired every 1 second for the duration of the experiment.

### 3D Strain Quantification and Imaris Strain Mapping

For the image segmentation and strain quantification of the NMBS in 3D we developed an Imaris (Bitplane) and MATLAB (Mathworks) image analysis pipeline. 3D image stacks obtained from confocal fluorescence microscopy were imported into Imaris for rendering and visualization. To identify intersection nodes of the NMBS, we first created a surface object of the NMBS based upon fluorescence intensity and performed a distance transformation. Nodes of the NMBS tend to have a high distance value when looking at the distance transformation within a surface. The Distance transformation was then used as the target image for spot detection in Imaris. Spots were placed at each NMBS intersection first through automated detection and then subsequent manual identification. Spot tracking through time insured that each node was properly identified and tracked throughout the entire time series. Once the spots were identified at each NMBS intersection node, a MATLAB script is run to initiate node to node pair identification and strain calculation.

Imaris Xtensions allow for transfer of data between Imaris and MATLAB for enhanced image analysis and quantification. Following node identification after spot creation and tracking, we developed a MATLAB-based Imaris Xtension to calculate 3D strain from the fluorescence NMBS images. Coordinates for each spot in X, Y, Z were exported and read into MATLAB. An image of the NMBS in its reference state was imported and maximally intensity projected into a 2D reference image for spot overlay. A skeletonized 2D version of the NMBS at time 0 was used as a road map for node pair assignment. We implemented an algorithm to iteratively traverse the skeletonized image and connect paired nodes. In the event of improper pair assignment, manual correction is made to either add or remove node pairs. Once all node pairs have been correctly assigned, the 3D distance between each pair is calculated and used for subsequent strain determination by calculating the (measured length / reference length) - 1.

We next created an import Xtension in Imaris to import each segment strain value as a filament object in Imaris that can be color coded with a strain statistic. Filaments were created for all node pairs and tracked for each time point. The change in segment length of the NMBS was then visualized as a color-coded strain statistic normalized around 0. Positive strain values corresponded to tensile strain and negative strain corresponded to compressive strain. Individual or groups of segments can be selected for additional data analysis and quantification as an exported Microsoft Excel data sheet. All MATLAB scripts and functions as well as Imaris Xtensions have been made freely available and open source through the Imaris Open File Exchange (http://open.bitplane.com/). Images of the NMBS strain at selected time points and videos showing 3D changes in strain throughout time can be exported from Imaris using the built-in animation capabilities.

### FFT and Beat Frequency Mapping

To begin, fluorescence images of spontaneously beating and electrically stimulated cardiomyocytes with 555-FN-NMBS applied were acquired via Micro-Manager software on an inverted Nikon Eclipse TI wide-field fluorescence microscope with a Nikon 20X Plan Apo Lambda objective, a Prime 95B Scientific CMOS camera (Photometrics), X-Cite 120PC light source (Excelitas Inc.), and appropriate excitation and emission filters for GFP and Tx-Red fluorescence. Cardiomyocytes were imaged in Tyrode’s medium containing 5 µM calcium indicator dye Cal 520 AM (21130, AAT Bioquest), and paced at 1 and 2 Hz using two parallel platinum electrodes with an 80V, 20 ms, square wave pulse applied with a Grass Stimulator. Time series images were imported into MATLAB software for further image analysis.

A custom MATLAB code (BeatFrequencyFFTMapping.m) was created to determine and evaluate the heterogeneity in the beat frequency across a field of cardiomyocytes from the NMBS motion and the fluctuations in calcium indicator dye fluorescence intensity. Time series images were first downsampled and gaussian blurred to decrease the effects of noise. To extract the beat frequency information from the fluorescence images, fluorescence intensity over time was measured for each pixel within the image series. The time series data of fluorescence intensity was then smoothed using a Savitsky-Golay filter. A Fast Fourier Transform (FFT) of the smoothed fluorescence intensity over time was performed to extract the dominant frequency for each pixel location from the peak of the power spectra density plot^58^. A pseudo image was then created consisting of the dominant frequency at each pixel location within the image and a JET look up table was utilized to visualize beat frequency heterogeneity. To regain the original image resolution the pseudo image was upsampled resulting in a final frequency map (Figure 4d). For the NMBS FFT analysis the final frequency map was pass through filtered with a binary mask of the original NMBS fluorescence intensity to restrict the analysis to the boundaries of the time 0 NMBS location.

### Drosophila Ovariole Dissection and Isolation

Fly stocks used in this study: *Moesin-GFP/TM6,Tb*^59,60^ and *P{His2Av-mRFP1}III.1* (Blooming Drosophila Stock Center #23650). A fly stock containing both Moesin-GFP and H2Av-RFP was generated by crossing the aforementioned parental fly lines (*P{His2Av-mRFP1}III.1, Moesin-GFP/TM6,Tb)*. All fly cultures were reared, and adult progeny maintained at 22-23°C/70% relative humidity/12-12hr light-dark cycle. One to two-week old adult female flies expressing either Moesin-GFP or a combination of Moesin-GFP and Histone 2A-RFP were transferred to vials containing cornmeal fly food and yeast paste for five days. On the fifth day, ovaries were extracted from the female flies in 22°C unsupplemented Grace’s media (Thermo, Cat#: 11595030) using standard Dumont forceps. For each set of ovaries, the outer fibrous sheath was removed and individual ovarioles were teased apart. Teased ovaries were kept in 22°C unsupplemented Grace’s media until ready for mesh application on the same day.

### Statistics and Data Analysis

Statistical and graphical analyses were performed using GraphPad Prism 7 software. Statistical tests were chosen based on the experimental sample size, distribution, and data requirements. For statistical analysis of the uniaxial tensile testing results data was plotted as mean ± standard deviation (S.D.) and evaluated for correlation using a Pearson correlation coefficient. For quantification and evaluation of NMBS segmental strain, individual strain segments of interest were selected in Imaris and exported for analysis in Excel (Microsoft). Identified regions showing similar tensile or compressive strain were averaged and plotting in Prism 7 (GraphPad) as mean ± standard deviation. Preparation of figures and visuals were constructed in Adobe Photoshop and Illustrator CS6. Fluorescence images were edited in Fiji (ImageJ NIH) and Imaris 9.1.2 (Bitplane). Advanced image analysis and quantification was performed in MATLAB (Mathworks) using custom software as mentioned previously.

## Acknowledgements

We thank M. Puthenveedu for use of the spinning disk confocal microscope; T. Cohen-Karni for use of upright resonant scanning confocal microscope; T. Hinton for assistance with 3D printing and laser-cutting, and R. Palchesko for SIA and microfabrication assistance. Research reported in this publication was supported by grants from the Human Frontier Science Program (RGY0071) and National Institutes of Health (DP2HL117750), and the National Heart, Lung, And Blood Institute under Award Number F32HL142229.

## Author contributions

All authors conceived the experiments and contributed to the scientific planning and discussions. D.J.S. prepared final figures and text. D.J.S. conducted all live confocal imaging of C2C12, and HSMC with NMBS. D.J.S. and M.B conducted Drosophila experiments and analysis. D.J.S. and J.T. wrote custom analysis code in MATLAB and Imaris. A.T. conducted tensile testing experiments and analysis. J.B. performed cardiomyocyte stem cell culture. E.A. and A.T. performed FEA simulation and analysis. J.M.S. and Q.J. assisted with the uniaxial tensile testing and wide-field fluorescence microscopy. D.J.S., J.T., A.T., E.A., J.B., B.M., and A.W.F. wrote the manuscript and interpreted the data.

## References

1 Bleaken, B. M., Menko, A. S. & Walker, J. L. Cells activated for wound repair have the potential to direct collective invasion of an epithelium. Mol. Biol. Cell 27, 451–465, doi:10.1091/mbc.E15-09-0615 (2015).

2 Boucher, E. & Mandato, C. A. Plasma membrane and cytoskeleton dynamics during single-cell wound healing. Biochim. Biophys. Acta 1853, 2649–2661, doi:10.1016/j.bbamcr.2015.07.012 (2015).

3 Rolin, G. L. et al. In vitro study of the impact of mechanical tension on the dermal fibroblast phenotype in the context of skin wound healing. J. Biomech. 47, 3555–3561, doi:10.1016/j.jbiomech.2014.07.015 (2014).

4 Basan, M., Elgeti, J., Hannezo, E., Rappel, W. J. & Levine, H. Alignment of cellular motility forces with tissue flow as a mechanism for efficient wound healing. Proc. Natl Acad. Sci. USA 110, 2452–2459, doi:10.1073/pnas.1219937110 (2013).

5 Wiegand, C. & White, R. Microdeformation in wound healing. Wound Repair Regen. 21, 793–799, doi:10.1111/wrr.12111 (2013).

6 Valero, C., Javierre, E., Garcia-Aznar, J. M. & Gomez-Benito, M. J. A cell-regulatory mechanism involving feedback between contraction and tissue formation guides wound healing progression. PloS one 9, e92774, doi:10.1371/journal.pone.0092774 (2014).

7 Kim, Y. et al. Mechanochemical actuators of embryonic epithelial contractility. Proceedings of the National Academy of Sciences of the United States of America 111, 14366–14371, doi:10.1073/pnas.1405209111 (2014).

8 Munjal, A., Philippe, J. M., Munro, E. & Lecuit, T. A self-organized biomechanical network drives shape changes during tissue morphogenesis. Nature 524, 351–355, doi:10.1038/nature14603 (2015).

9 Kiehart, D. P. Epithelial morphogenesis: apoptotic forces drive cell shape changes. Dev. Cell 32, 532–533, doi:10.1016/j.devcel.2015.02.020 (2015).

10 Petridou, N. I. & Skourides, P. A. FAK transduces extracellular forces that orient the mitotic spindle and control tissue morphogenesis. Nat. Commun. 5, 5240, doi:10.1038/ncomms6240 (2014).

11 Martin, A. C. & Goldstein, B. Apical constriction: themes and variations on a cellular mechanism driving morphogenesis. Development 141, 1987–1998, doi:10.1242/dev.102228 (2014).

12 Munjal, A. & Lecuit, T. Actomyosin networks and tissue morphogenesis. Development 141, 1789–1793, doi:10.1242/dev.091645 (2014).

13 Heller, E., Kumar, K. V., Grill, S. W. & Fuchs, E. Forces generated by cell intercalation tow epidermal sheets in mammalian tissue morphogenesis. Dev. Cell 28, 617–632, doi:10.1016/j.devcel.2014.02.011 (2014).

14 Aguilar-Cuenca, R., Juanes-Garcia, A. & Vicente-Manzanares, M. Myosin II in mechanotransduction: master and commander of cell migration, morphogenesis, and cancer. Cell. Mol. Life Sci. 71, 479–492, doi:10.1007/s00018-013-1439-5 (2014).

15 Paez-Mayorga, J. et al. Bioreactors for Cardiac Tissue Engineering. Adv Healthc Mater 8, doi:ARTN 1701504 10.1002/adhm.201701504 (2019).

16 Freytes, D. O., Wan, L. Q. & Vunjak-Novakovic, G. Geometry and force control of cell function. J Cell Biochem 108, 1047–1058, doi:10.1002/jcb.22355 (2009).

17 Liu, W. F. Mechanical regulation of cellular phenotype: implications for vascular tissue regeneration. Cardiovasc Res 95, 215–222, doi:10.1093/cvr/cvs168 (2012).

18 Hasenfuss, G. et al. Alteration of contractile function and excitation-contraction coupling in dilated cardiomyopathy. Circ Res 70, 1225–1232, doi:10.1161/01.res.70.6.1225 (1992).

19 Tschumperlin, D. J., Ligresti, G., Hilscher, M. B. & Shah, V. H. Mechanosensing and fibrosis. J Clin Invest 128, 74–84, doi:10.1172/JCI93561 (2018).

20 Navis, A. & Nelson, C. M. Pulling together: Tissue-generated forces that drive lumen morphogenesis. Semin. Cell Dev. Biol., doi: 10.1016/j.semcdb.2016.1001.1002, doi:10.1016/j.semcdb.2016.01.002 (2016).

21 Campas, O. et al. Quantifying cell-generated mechanical forces within living embryonic tissues. Nat. Methods 11, 183–189, doi:10.1038/nmeth.2761 (2014).

22 Ramasubramanian, A. & Taber, L. A. Computational modeling of morphogenesis regulated by mechanical feedback. Biomech Model Mechanobiol 7, 77–91, doi:10.1007/s10237-007-0077-y (2008).

23 Jackson, T. R., Kim, H. Y., Balakrishnan, U. L., Stuckenholz, C. & Davidson, L. A. Spatiotemporally Controlled Mechanical Cues Drive Progenitor Mesenchymal-to-Epithelial Transition Enabling Proper Heart Formation and Function. Curr Biol 27, 1326–1335, doi:10.1016/j.cub.2017.03.065 (2017).

24 Cai, D. et al. Mechanical feedback through E-cadherin promotes direction sensing during collective cell migration. Cell 157, 1146–1159, doi:10.1016/j.cell.2014.03.045 (2014).

25 Munevar, S., Wang, Y. & Dembo, M. Traction force microscopy of migrating normal and H-ras transformed 3T3 fibroblasts. Biophys J 80, 1744–1757, doi:10.1016/s0006-3495(01)76145-0 (2001).

26 Fu, J. et al. Mechanical regulation of cell function with geometrically modulated elastomeric substrates. Nature methods 7, 733–736, doi:10.1038/nmeth.1487 (2010).

27 Legant, W. R. et al. Multidimensional traction force microscopy reveals out-of-plane rotational moments about focal adhesions. Proceedings of the National Academy of Sciences of the United States of America 110, 881–886, doi:10.1073/pnas.1207997110 (2013).

28 Schluck, T. & Aegerter, C. M. Photo-elastic properties of the wing imaginal disc of Drosophila. Eur. Phys. J. E Soft Matter 33, 111–115, doi:10.1140/epje/i2010-10580-8 (2010).

29 Sugimura, K. & Ishihara, S. The mechanical anisotropy in a tissue promotes ordering in hexagonal cell packing. Development 140, 4091–4101, doi:10.1242/dev.094060 (2013).

30 Sun, Y., Jallerat, Q., Szymanski, J. M. & Feinberg, A. W. Conformal nanopatterning of extracellular matrix proteins onto topographically complex surfaces. Nature methods 12, 134–136, doi:10.1038/nmeth.3210 (2015).

31 Feinberg, A. W. & Parker, K. K. Surface-initiated assembly of protein nanofabrics. Nano letters 10, 2184–2191, doi:10.1021/nl100998p (2010).

32 Palchesko, R. N., Szymanski, J. M., Sahu, A. & Feinberg, A. W. Shrink Wrapping Cells in a Defined Extracellular Matrix to Modulate the Chemo-Mechanical Microenvironment. Cell Mol Bioeng 7, 355–368, doi:10.1007/s12195-014-0348-5 (2014).

33 Palchesko, R. N., Funderburgh, J. L. & Feinberg, A. W. Engineered Basement Membranes for Regenerating the Corneal Endothelium. Adv Healthc Mater 5, 2942–2950, doi:10.1002/adhm.201600488 (2016).

34 Nunes, L. C. S. Mechanical characterization of hyperelastic polydimethylsiloxane by simple shear test. Mat Sci Eng a-Struct 528, 1799–1804, doi:10.1016/j.msea.2010.11.025 (2011).

35 Brown, A. E., Litvinov, R. I., Discher, D. E., Purohit, P. K. & Weisel, J. W. Multiscale mechanics of fibrin polymer: gel stretching with protein unfolding and loss of water. Science 325, 741–744, doi:10.1126/science.1172484 (2009).

36 Schwarz, U. S. & Soine, J. R. Traction force microscopy on soft elastic substrates: A guide to recent computational advances. Biochimica et biophysica acta 1853, 3095–3104, doi:10.1016/j.bbamcr.2015.05.028 (2015).

37 Dembo, M. & Wang, Y. L. Stresses at the cell-to-substrate interface during locomotion of fibroblasts. Biophysical Journal 76, 2307–2316, doi:Doi 10.1016/S0006-3495(99)77386-8 (1999).

38 Stapleton, S. C., Chopra, A. & Chen, C. S. Force measurement tools to explore cadherin mechanotransduction. Cell Commun Adhes 21, 193–205, doi:10.3109/15419061.2014.905929 (2014).

39 Bers, D. M. Cardiac excitation-contraction coupling. Nature 415, 198–205, doi:10.1038/415198a (2002).

40 Jacot, J. G., Martin, J. C. & Hunt, D. L. Mechanobiology of cardiomyocyte development. Journal of biomechanics 43, 93–98, doi:10.1016/j.jbiomech.2009.09.014 (2010).

41 Ribeiro, M. C. et al. Functional maturation of human pluripotent stem cell derived cardiomyocytes in vitro--correlation between contraction force and electrophysiology. Biomaterials 51, 138–150, doi:10.1016/j.biomaterials.2015.01.067 (2015).

42 Andersen, D. & Horne-Badovinac, S. Influence of ovarian muscle contraction and oocyte growth on egg chamber elongation in Drosophila. Development 143, 1375–1387, doi:10.1242/dev.131276 (2016).

43 Roth, S. & Lynch, J. A. Symmetry breaking during Drosophila oogenesis. Cold Spring Harb Perspect Biol 1, a001891, doi:10.1101/cshperspect.a001891 (2009).

44 Pearson, J. R. et al. ECM-Regulator timp Is Required for Stem Cell Niche Organization and Cyst Production in the Drosophila Ovary. PLoS Genet 12, e1005763, doi:10.1371/journal.pgen.1005763 (2016).

45 Palchesko, R. N., Zhang, L., Sun, Y. & Feinberg, A. W. Development of polydimethylsiloxane substrates with tunable elastic modulus to study cell mechanobiology in muscle and nerve. PloS one 7, e51499, doi:10.1371/journal.pone.0051499 (2012).

46 Klotzsch, E. et al. Fibronectin forms the most extensible biological fibers displaying switchable force-exposed cryptic binding sites. Proceedings of the National Academy of Sciences of the United States of America 106, 18267–18272, doi:10.1073/pnas.0907518106 (2009).

47 Muiznieks, L. D. & Keeley, F. W. Molecular assembly and mechanical properties of the extracellular matrix: A fibrous protein perspective. Biochimica et biophysica acta 1832, 866–875, doi:10.1016/j.bbadis.2012.11.022 (2013).

48 Argudo, D., Bethel, N. P., Marcoline, F. V., Wolgemuth, C. W. & Grabe, M. New Continuum Approaches for Determining Protein-Induced Membrane Deformations. Biophys J 112, 2159–2172, doi:10.1016/j.bpj.2017.03.040 (2017).

49 Argudo, D., Bethel, N. P., Marcoline, F. V. & Grabe, M. Continuum descriptions of membranes and their interaction with proteins: Towards chemically accurate models. Biochimica et biophysica acta 1858, 1619–1634, doi:10.1016/j.bbamem.2016.02.003 (2016).

50 Yoshihiro Ujihara, M. N., and Shigeo Wada. A Mechanical Cell Model and Its Application to Cellular Biomechanics. IntechOpen (2011).

51 Szymanski, J. M., Jallerat, Q. & Feinberg, A. W. ECM protein nanofibers and nanostructures engineered using surface-initiated assembly. J. Vis. Exp., doi: 10.3791/51176, doi:10.3791/51176 (2014).

52 Szymanski, J. M. & Feinberg, A. W. Fabrication of freestanding alginate microfibers and microstructures for tissue engineering applications. Biofabrication 6, 024104, doi:10.1088/1758-5082/6/2/024104 (2014).

53 Duffy, R. M., Sun, Y. & Feinberg, A. W. Understanding the Role of ECM Protein Composition and Geometric Micropatterning for Engineering Human Skeletal Muscle. Ann Biomed Eng 44, 2076–2089, doi:10.1007/s10439-016-1592-8 (2016).

54 Lian, X. et al. Directed cardiomyocyte differentiation from human pluripotent stem cells by modulating Wnt/beta-catenin signaling under fully defined conditions. Nat Protoc 8, 162–175, doi:10.1038/nprot.2012.150 (2013).

55 Lee, A. et al. 3D bioprinting of collagen to rebuild components of the human heart. Science 365, 482–487, doi:10.1126/science.aav9051 (2019).

56 Xu, X. Q., Soo, S. Y., Sun, W. & Zweigerdt, R. Global expression profile of highly enriched cardiomyocytes derived from human embryonic stem cells. Stem Cells 27, 2163–2174, doi:10.1002/stem.166 (2009).

57 Burridge, P. W. et al. Chemically defined generation of human cardiomyocytes. Nature methods 11, 855–860, doi:10.1038/nmeth.2999 (2014).

58 Reno, A., Hunter, A., Li, Y., Ye, T. & Foley, A. Quantification of Cardiomyocyte Beating Frequency Using Fourier Transform Analysis. Photonics 5, doi:10.3390/photonics5040039 (2018).

59 Edwards, R. G. & Beard, H. K. Oocyte polarity and cell determination in early mammalian embryos. Mol Hum Reprod 3, 863–905, doi:10.1093/molehr/3.10.863 (1997).

60 Edwards, K. A., Demsky, M., Montague, R. A., Weymouth, N. & Kiehart, D. P. GFP-moesin illuminates actin cytoskeleton dynamics in living tissue and demonstrates cell shape changes during morphogenesis in Drosophila. Dev Biol 191, 103–117, doi:10.1006/dbio.1997.8707 (1997).

