## Supplemental Figures for "yFibronectin-Based Nanomechanical Biosensors to Map 3D Strains in Live Cells and Tissues"

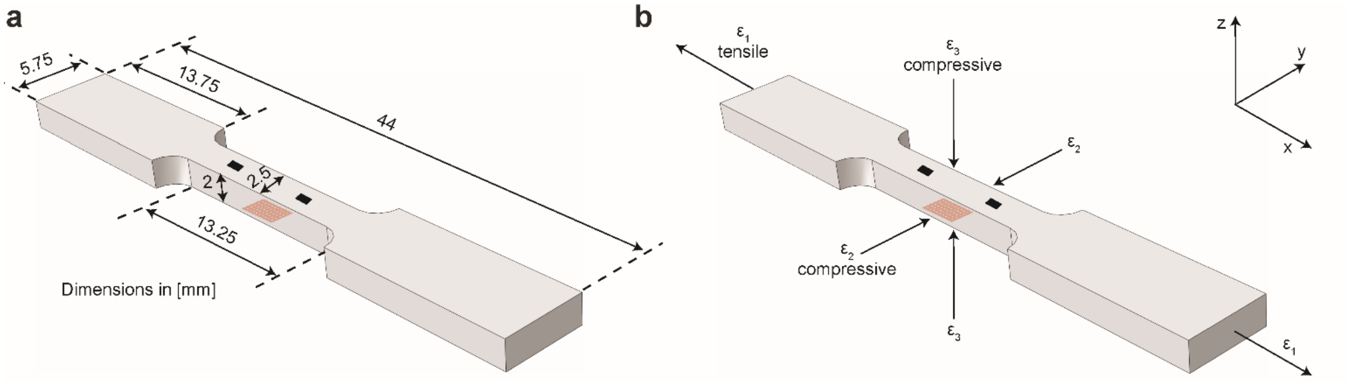


**Figure S1 | Design and dimensions of tensile testing strip. a.** The overall design of the tensile testing strip was chosen to mimic a classic "dog bone" structure commonly used for uniaxial mechanical testing. The NMBS was applied to the underneath side of the hydrogel test strip and macro fiduciary marks were applied to the top surface for tracking during tensile testing. **b.** Mechanical strains were defined as $\varepsilon_1$ (tensile, X, in the direction of elongation), $\varepsilon_2$ (compressive, Y, perpendicular to the direction of elongation), and $\varepsilon_3$ (compressive, Z, orthogonal to the XY axis).

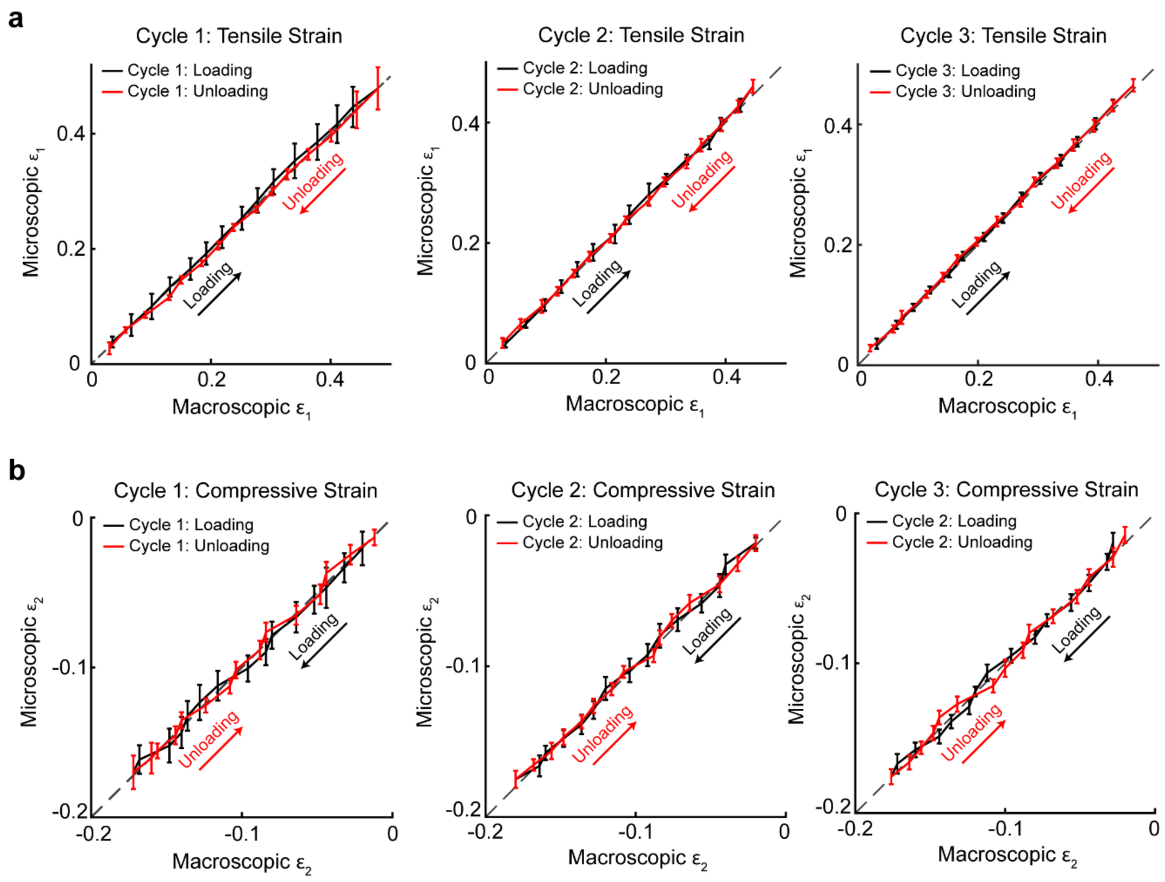


**Figure S2 | Measuring microscopic PDMS strain with the NMBS during uniaxial cyclic loading. a.** Results from three cycles of cyclic loading showing accurate NMBS tensile strain $\varepsilon_1$ tracking during loading and unloading (mean ± S.D.). **b.** Results from three cycles of cyclic loading showing accurate NMBS compressive strain $\varepsilon_2$ tracking during loading and unloading (mean ± S.D.).

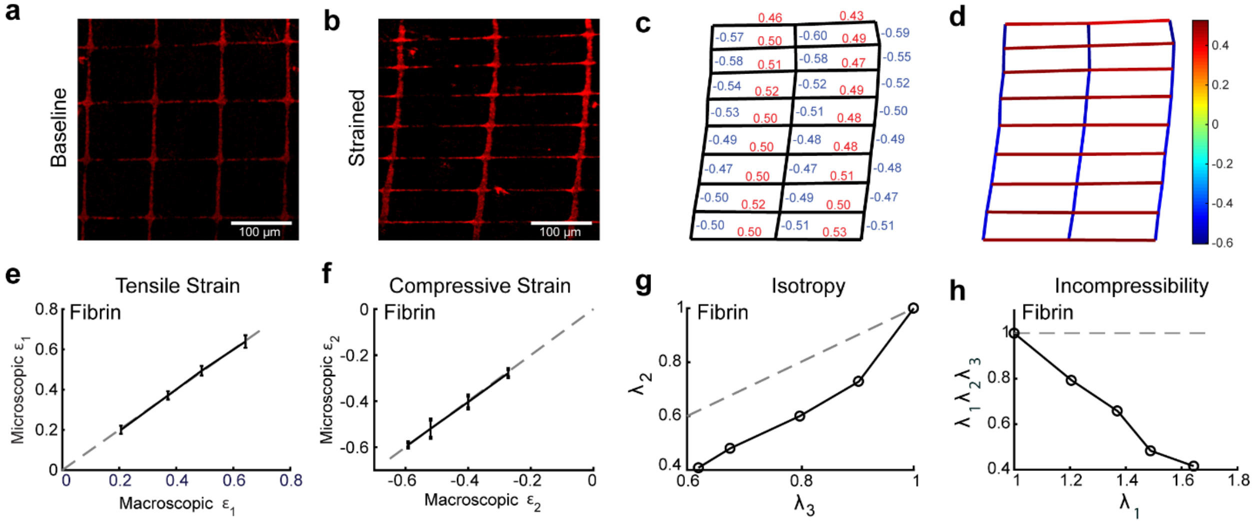


**Figure S3 | Uniaxial tensile testing and Finite Element Analysis of the NMBS on a Fibrin test strip.** **a.** Example wide-field fluorescence image of the pre-strain NMBS (100 µm X 100 µm X 10 µm) on a fibrin test strip. **b**. Representative fluorescence image of 50% strained NMBS following tensile testing. **c**. Image segmentation and analysis of NMBS nodes and segments following 50% uniaxial tensile (positive) and compressive (negative) mechanical strain. **d**. A color map indicating the locations and magnitude of tensile and compressive mechanical strains. **e.** The microscopic tensile strain $\varepsilon_1$ of the NMBS strongly correlates with the macroscopic tensile strain $\varepsilon_1$ of fibrin (mean ± S.D.). **f**. The microscopic compressive strain $\varepsilon_2$ of the NMBS strongly correlates with the macroscopic compressive strain $\varepsilon_2$ of fibrin (mean ± S.D.). **g.** For fibrin gels the There stretch ratios $\lambda_2$ and $\lambda_3$ were not equivalent confirming that fibrin is an anisotropic material. **h.** The product of the stretch ratios $\lambda_1\lambda_2\lambda_3$ was less than 1 during uniaxial stretching confirming that fibrin is compressible.

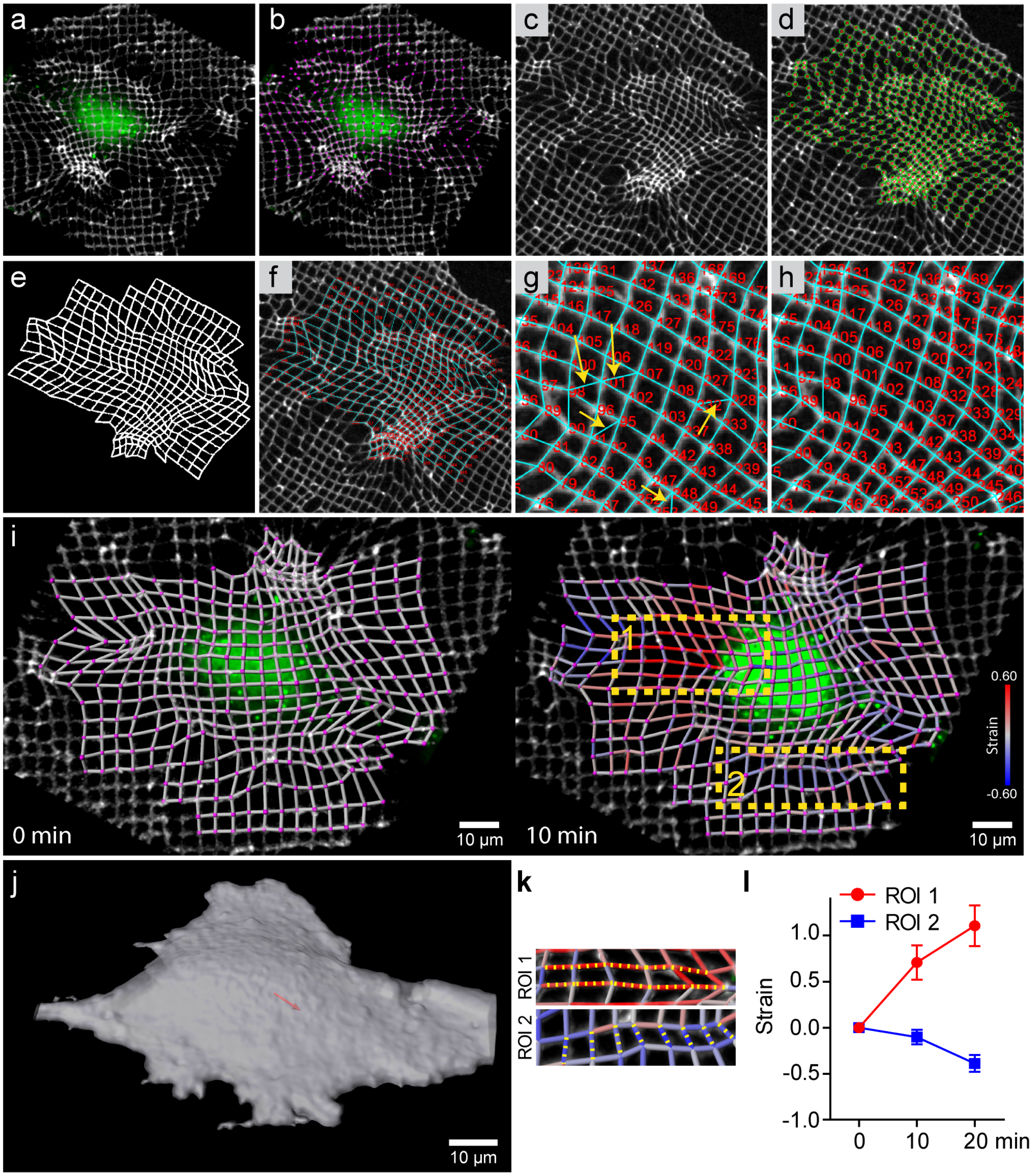


**Figure S4 | Custom Imaris and Matlab segmentation and tracking of NMBS images for strain quantification. a.** Confocal fluorescence image of 633-FN-NMBS (2 μm X 2 μm X 2 μm, Gray) applied to HSMC incubated with 488-CellTracker (Green) and visualized in Imaris 3D visualization software. **b.** NMBS mesh node detection (Magenta) overlaid onto NMBS. **c.** Maximum intensity Z-projection of NMBS imported into Matlab from Imaris. **d.** X and Y node coordinates (Green circles) exported from

Imaris to Matlab and overlaid onto NMBS maximum intensity Z-projection for pair determination. **e.** Skeletonized binary image of the NMBS mesh defined by the node coordinates. **f.** Computationally generated pair determination via pathfinding algorithm. **g.** Example errors in pair determination (yellow arrows). Numbers represent the assigned i.d. for each individual paired line segment. **h.** Manual correction of wrongly assigned pair-wise connections. **i.** Paired segments between connected nodes imported from Matlab to Imaris with a custom statistic for mapping engineering strain (baseline strain = 0, gray; tension = positive values, red; compression = negative values, blue). **j.** Displacement vector (red arrow) following cell tracking showing direction of HSMC migration during imaging sequence. **k.** Two regions of interest (ROI 1, tensile region; ROI 2, compressive region) extracted for quantitative strain analysis from "i". **l.** ROI 1 displayed increased strain while ROI 2 showed decreasing strain as the cell migrated to the right of the image (mean ± S.D.).

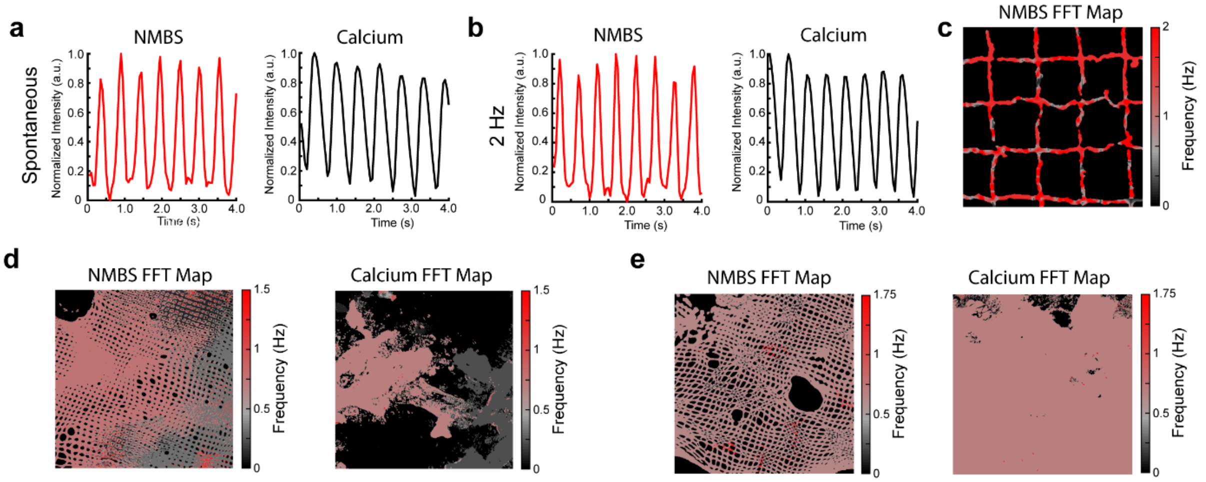


**Figure S5 | Utilizing the NMBS to track functional cardiomyocyte performance. a.** Pixel level changes in fluorescence intensity for NMBS and calcium (Fluo-4 dye) during spontaneous iPSC-derived cardiomyocyte contractions. **b.** Changes in fluorescence intensity for NMBS and calcium (Fluo-4 dye) during 2 Hz field stimulation of iPSC-derived cardiomyocytes. **c.** Fast Fourier Transform (FFT) map of predominant cardiomyocyte beat frequency obtained from NMBS (100 µm X 100 µm X 10 µm) fluorescence motion following 2 Hz field stimulation. **d.** Heterogeneous FFT map from NMBS (20 µm X 20 µm X 10 µm) and calcium (Fluo-4 dye) of spontaneously contracting iPSC-derived cardiomyocytes. **e.** Homogeneous FFT map from NMBS (20 µm X 20 µm X 10 µm) and calcium (Fluo-4 dye) of spontaneously contracting iPSC-derived cardiomyocytes.

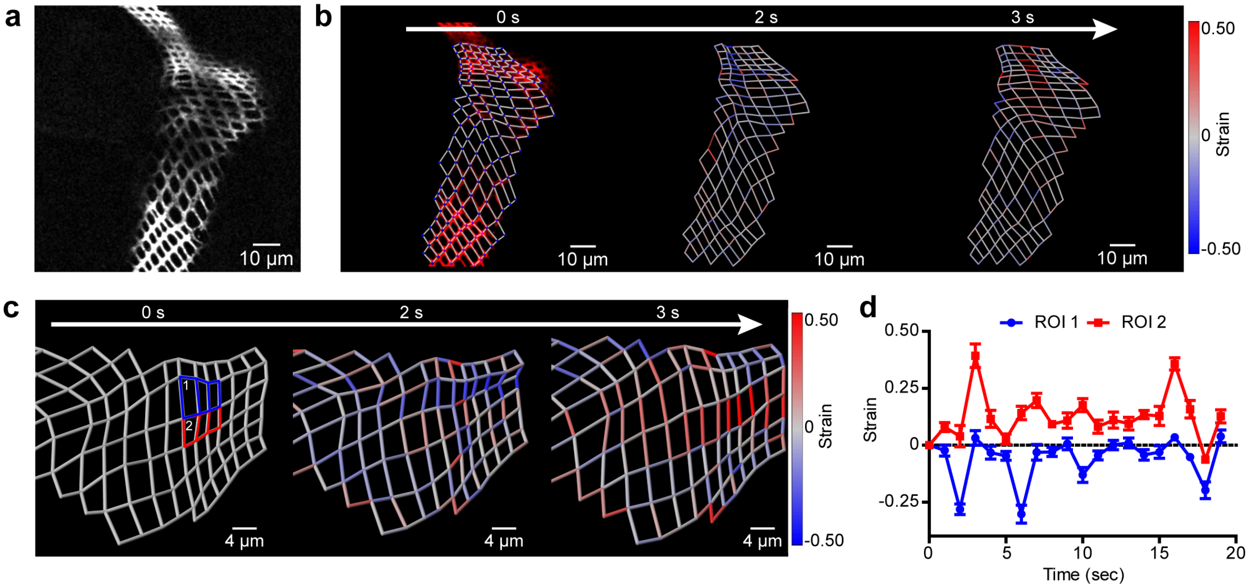


**Figure S6 | Live strain via NMBS mapping during ovariole muscle sheath contractions. a.** Maximum intensity Z projection of spinning disk confocal fluorescence images of a region of NMBS applied to a drosophila ovariole. **b.** Overlay of segmented and tracked NMBS (2 μm X 2 μm X 2 μm) segments and nodes onto the NMBS fluorescence image. Tension (red) and compression (blue) are regionally localized during ovariole muscle contractions. **c.** Region of interest (ROI 1 (blue), and 2 (red)) that undergoes large compressive and tensile strain over time. **d.** Quantification of ROI 1 and 2 showing oscillations in strain during ovariole contractile motion (mean ± S.D.).
